# Impact of SNP microarray analysis of compromised DNA on kinship classification success in the context of investigative genetic genealogy

**DOI:** 10.1101/2021.06.25.449870

**Authors:** Jard H. de Vries, Daniel Kling, Athina Vidaki, Pascal Arp, Vivian Kalamara, Michael M.P.J. Verbiest, Danuta Piniewska-Róg, Thomas J. Parsons, André G. Uitterlinden, Manfred Kayser

## Abstract

Single nucleotide polymorphism (SNP) data generated with microarray technologies have been used to solve murder cases via investigative leads obtained from identifying relatives of the unknown perpetrator included in accessible genomic databases, referred to as investigative genetic genealogy (IGG). However, SNP microarrays were developed for relatively high input DNA quantity and quality, while SNP microarray data from compromised DNA typically obtainable from crime scene stains are largely missing. By applying the Illumina Global Screening Array (GSA) to 264 DNA samples with systematically altered quantity and quality, we empirically tested the impact of SNP microarray analysis of deprecated DNA on kinship classification success, as relevant in IGG. Reference data from manufacturer-recommended input DNA quality and quantity were used to estimate genotype accuracy in the compromised DNA samples and for simulating data of different degree relatives. Although stepwise decrease of input DNA amount from 200 nanogram to 6.25 picogram led to decreased SNP call rates and increased genotyping errors, kinship classification success did not decrease down to 250 picogram for siblings and 1^st^ cousins, 1 nanogram for 2^nd^ cousins, while at 25 picogram and below kinship classification success was zero. Stepwise decrease of input DNA quality via increased DNA fragmentation resulted in the decrease of genotyping accuracy as well as kinship classification success, which went down to zero at the average DNA fragment size of 150 base pairs. Combining decreased DNA quantity and quality in mock casework and skeletal samples further highlighted possibilities and limitations. Overall, GSA analysis achieved maximal kinship classification success from 800-200 times lower input DNA quantities than manufacturer-recommended, although DNA quality plays a key role too, while compromised DNA produced false negative kinship classifications rather than false positive ones.

**Author Summary:** Investigative genetic genealogy (IGG), i.e., identifying unknown perpetrators of crime via genomic database-tracing of their relatives by means of microarray-based single nucleotide polymorphism (SNP) data, is a recently emerging field. However, SNP microarrays were developed for much higher DNA quantity and quality than typically available from crime scenes, while SNP microarray data on quality and quantity compromised DNA are largely missing. As first attempt to investigate how SNP microarray analysis of quantity and quality compromised DNA impacts kinship classification success in the context of IGG, we performed systematic SNP microarray analyses on DNA samples below the manufacturer-recommended quantity and quality as well as on mock casework samples and on skeletal remains. In addition to IGG, our results are also relevant for any SNP microarray analysis of compromised DNA, such as for the DNA prediction of appearance and biogeographic ancestry in forensics and anthropology and for other purposes.

## Introduction

For almost three decades, forensic DNA profiling with standard sets of polymorphic short tandem repeat (STR) markers has successfully been used to identify perpetrators of crime, thereby contributing towards solving numerous criminal cases worldwide (1, 2). However, in principle, forensic STR profiling is unsuitable for identifying unknown perpetrators, whose STR profiles are not yet included in national forensic DNA databases or are unknown to the investigative authorities otherwise. This consequently leads to cold cases with STR profiles of crime scene stains being available but not matching any known suspect, including all criminal offenders stored in the national forensic DNA database. Such situation allows perpetrators to continue their criminal activities and justice is denied to victims of crime or their families.

Over the last years, several DNA-based approaches for tracing unknown perpetrators have emerged. One such way is to search for family members of the unknown stain donor based on the forensic STR-profiles in the national forensic DNA database (i.e., familial search) (3). This approach, however, is limited in its success to first degree relatives, because of the limited number of autosomal STRs used in routine forensic DNA profiling (4). This disadvantage can be overcome by applying male-specific STRs from non-recombining regions of the Y-chromosome (Y-STRs) that can highlight male relatives of the unknown male perpetrator, given that the vast majority of perpetrators of major crimes are males (5). However, because forensic DNA databases in almost all countries consist of autosomal STR profiles, but not Y-STR profiles, Y-STR based familial search is restricted to voluntary DNA mass screenings (5). A more indirect way to find unknown perpetrators via focused police investigation is through investigative leads obtained via the prediction of externally visible characteristics of the unknown stain donor from crime scene DNA, including appearance traits (6), bio-geographic ancestry (7), and chronological age (8), in the context of Forensic DNA Phenotyping (9). Most recently, investigative genetic genealogy (IGG) has started to emerge as new approach to find unknown perpetrators with the help of DNA (4, 10).

IGG, also known as forensic genetic genealogy (FGG), is based on genomic data from hundreds of thousands of autosomal single nucleotide polymorphisms (SNPs) typically generated with SNP microarray technology (10). Because of the large number of autosomal SNPs involved, IGG allows close and distant relatives from both, maternal and paternal sides to be identified (4, 11). Over the last years, genomic databases consisting of high-density SNP data have emerged, albeit outside the forensic field. Instead of governmental authorities, these genetic genealogy databases are managed by private companies or private citizens, such as GEDmatch consisting of around 1.3 million high-density SNP profiles as of 2020 (12). The most commonly used method for identifying relatives in IGG is via DNA segments shared between the unknown perpetrator obtained from crime scene DNA and individuals in genomic databases (10, 13, 14). These identical-by-descend (IBD) segments that originate from the same ancestor signal a familial relation between the highlighted person in the genomic database and the unknown perpetrator, and the length of the shared segments translates into how close the family relationship is. Police investigation to find the unknown perpetrator is then focused on the highlighted relative via genealogical research (10).

The recent resolution of the Golden State Killer case in the USA, together with several other cases, demonstrated the power of IGG (15, 16). As of November 2020, IGG had assisted in over 200 cold cases, of which at least 28 were solved (17) (18). In May 2019, GEDmatch, the most prominent genomic database used for IGG, updated its privacy regulations to require users to opt-in for law enforcement to search their SNP profiles, and by October 2019, only ∼185,000 GEDmatch users had done this (12), reflecting a dramatic decrease of law enforcement access compared to previous times when GEDmatch access for law enforcement was unrestricted. In December 2019, the GEDmatch genomic database was acquired by the forensic genomics firm Verogen. The company FamilyTreeDNA (FTDNA) also hosts a SNP microarray database of 2 million of its customers that was initially established for other than forensic reasons, while the company actively works with law enforcement to permit database access for IGG (12, 19). FTDNA customers now need to opt-out to restrict law enforcement from using their SNP profiles for searches. Currently, all genomic databases available for IGG consist of SNP microarray data obtained from DNA of customer collected cheek swab or saliva samples, also known as direct-to-consumer genetic testing (10). In principle, SNP profiles extracted from whole genome sequencing (WGS) data obtained from crime scene DNA can also be used to search DTC SNP databases given the SNP overlap, which was recently exemplified in a murder case in Sweden (20). However, genomic databases for IGG that are based on WGS data are yet to be established.

Besides the various ethical, societal, regulatory and other dimensions in relation to the forensic use of genomic databases that were not established for forensic purposes (4, 10, 21), there is an important technical dimension related to SNP microarray typing of forensic DNA such as for IGG purposes. Notably, all currently available SNP microarrays were developed and optimized for relatively high input DNA quality and quantity, which typically is not available from human biological stains found at crime scenes. Moreover, the DNA hybridization principle underlying all SNP microarray technologies is not expected to be well-suited for compromised input DNA of low quantity and quality typically available from crime scene stains. While the statistical methods to genetically classify relatives from high-density SNP data in the context of IGG are being established throughout the last years (14, 22), studies that applied SNP microarrays to compromised DNA samples are scarce (23–25). More importantly, as far as we are aware, in-depth studies to systematically test for the impact the use of SNP microarrays in compromised DNA has on kinship classification success in the context of IGG are missing completely from the scientific literature as of yet. It has been recognized by several authors that the lack of empirical data on the performance of SNP microarrays in compromised DNA and its consequence on forensic use marks a serious problem for the forensic application of SNP microarrays in general and IGG in particular (4, 10, 26).

In this study, we performed systematic SNP microarray experiments on hundreds of DNA samples from multiple individuals with varying degrees of DNA input quantity and quality to test the impact of resulting SNP microarray genotyping errors on kinship classification success in the context of IGG. From the individual DNA samples of which we used DNA samples of varying quality and quantity for SNP microarray analysis, we also generated high-quality reference data from the high quality and quantity DNA input conditions recommended by the microarray manufacturer. Together, these data were used to calculate genotype accuracy for the compromised DNA samples. The high-quality data were additionally used to simulate data of different degree relatives for kinship classification. To get further insights, we additionally performed SNP microarray genotyping on DNA samples with a combined decrease of quantity and quality i.e., DNA from mock casework samples of varying storage temperature, time, and artificial degradation as well as naturally degraded DNA obtained from skeletal remains. The overall SNP microarray data set we generated in this study allowed us to quantify the effect of decreased DNA quality and quantity on SNP microarray genotyping quality with regard to the ability to classify relatives of different degrees of relationship. This data is vital for the development of IGG applications based on SNP microarray analysis of DNA obtained from human crime scene samples in forensic genetic casework, or from low quality and/or quantity DNA samples for other purposes in forensic casework and anthropological studies.

## Results

To evaluate SNP microarray technology for quantity and/or quality compromised input DNA and its consequence for kinship classification in the context of IGG, we performed a series of SNP microarray experiments with the widely used Illumina Global Screening Array (GSA). We started out by generating microarray data of 24 individuals by using high input DNA quantity and quality by meeting the recommendations of the microarray manufacturer i.e., 200 ng of high molecular weight DNA. This reference dataset was then used to conditionally simulate SNP data of relatives of these 24 individuals based on four degrees of relationships i.e., full siblings, 1^st^ cousins, 2^nd^ cousins, and 3^rd^ cousins (**Figure S1**). We also used this high-quality reference dataset to determine genotyping errors in the data obtained from the compromised DNA samples of the same individuals, respectively (see methods for details).

### Impact of decreased input DNA quantity on SNP microarray-based kinship classification

We quantified the impact of decreased input DNA quantity on the success of kinship classification based on SNP microarray data. For this, we generated 8 GSA datasets per each of 24 individuals based on manufacturer-recommended 200 ng input DNA as well as based on seven lower amounts i.e., 1000 pg, 250 pg, 100 pg, 50 pg, 25 pg, 12.5 pg, and 6.25 pg, and used them to perform kinship classification.

We found that on average the success of kinship classification remained over 98.5% for input DNA amounts from 200 ng down to 250 pg for siblings and 1^st^ cousins, and down to 1ng for 2^nd^ cousins (**Figure 1**). With 25 pg (approximately 4 cells worth of DNA) and lower input DNA amounts, the kinship classification success was zero for relatives of all four degrees (**Figure 1, Table S1**). We observed considerable variation between DNA samples of the same input amount between the 24 individuals on how decreasing input DNA quantity impacted on kinship classification success. Classification of siblings remained 100% correct for all 24 individuals from 200 ng down to 250 pg and for 17 individuals also with 100 pg (average classification rate of 93%). With 50 pg, 5 individuals still had 100% success rate for sibling classification, while with 25 pg 22 individuals showed 0% classification success for siblings and any other relatives tested. For 1^st^ cousins, 100% classification success was revealed for all 24 individuals with 200 ng and 1 ng, while with 250 pg it was for 22 individuals (average classification success of 99.4%), and with 100 pg for 13 individuals (76.6% success on average across individuals). None of the 24 individuals had 100% classification success when applying 50 pg input DNA (18.5% success on average). For 2^nd^ cousins, classification success was close to 99% for all 24 individuals with 200 ng, which remained stable down to 1 ng (98.6% success on average) and dropped to 91.6% with 250 pg, 55.7% with 100 pg and 6.6% with 50 pg. For 3^rd^ cousins, we first note that even with ideal input DNA quantity of the manufacturer-recommended 200 ng, classification success was on average only 74.6% across the 24 individuals, with no individual sample having more than 81.5% success rate, likely due to insufficient identical-by-descent (IBD) sharing. The classification success rate was slightly reduced from 1 ng input DNA (72.6% success). Using 250 pg input DNA, the 3^rd^ cousin classification success dropped down to 62.6%, with all but one sample decreasing in classification success by more than 1%, and further down to 30.7% with 100 pg, where all samples had decreased success rate but none yet at zero, and down to 3.3% with 50 pg, where 11 samples had 0% classification success. From input DNA amounts of 25 pg and lower, 3^rd^ cousin classification success was zero for all 24 individuals. When the true kinship relations were no longer classified correctly due to compromised input DNA quantity, misclassifications always occurred at the lower degrees of relatedness (e.g., siblings misclassified as cousins) and did not result in false overestimations of the classified degree of relationship.

**Figure 1:**
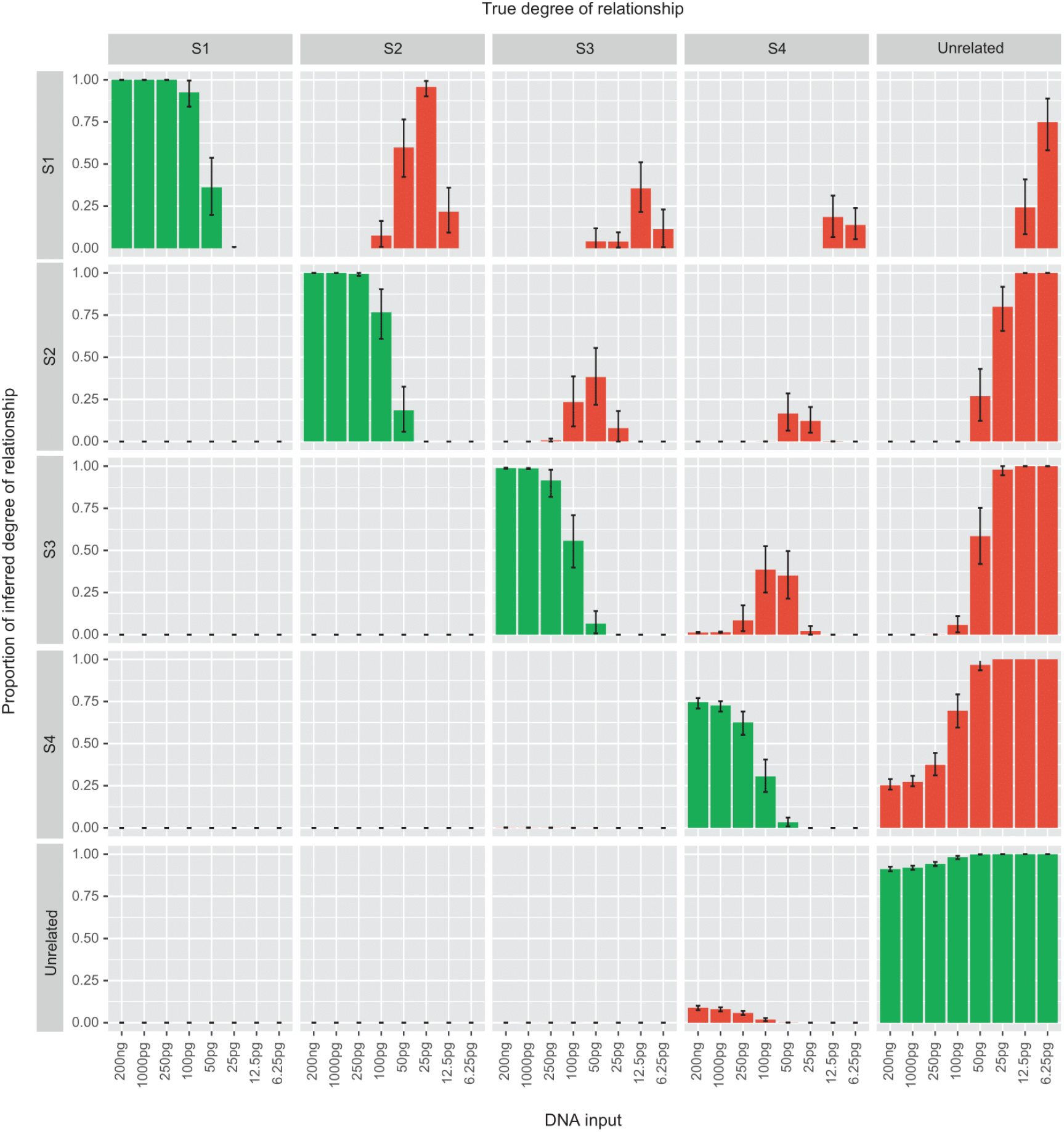
Kinship classification of simulated relatives based on SNP microarray genotype data obtained from input DNA of decreased quantity. Each plot in this matrix represents a simulated kinship relation and a specific kinship classification based on a SNP microarray dataset obtained from stepwise decreasing input DNA quantity. Each of the 24 individual DNA samples used in this experiment had 1000 relatives simulated for each degree of relative, which were S1 (Full siblings), S2 (1^st^ cousins), S3 (2^nd^ cousins), S4 (3^rd^ cousins) or unrelated from the high-quality reference data generated from manufacturer-recommended input-DNA quantity and quality; the kinship classification was restricted to these possible relation categories. For each of these 24 individuals, 8 input DNA quantity levels i.e., 200 ng, 1 ng, 250 pg, 100 pg, 50 pg, 25 pg, 12.5 pg, and 6.25 pg were processed on the GSA. The estimated genetic relation between such dataset from quantity-compromised DNA and the relatives simulated from high-quality reference data was then classified per each of the 24 individuals. The x-axis of the matrix describes the classified kinship relations, while the y-axis of the matrix describes the original simulated relations, across the 24 individuals used. For the individual plots, the x-axis describes the input DNA quantity, and the y-axis describes the average kinship classification rate. The green bar plots represent correctly classified relation and the red bar plots represent the incorrect classifications. Error bars display the 75% confidence interval.

### Effect of decreased input DNA quantity on SNP microarray genotype accuracy

Aiming to better understand the observed impact of decreased input DNA quantity on decreased kinship classification success, we investigated the effect of decreased DNA quantity on SNP microarray genotype accuracy by calculating microarray-based SNP call rates and genotype errors rates depending on varying input DNA amounts (as detailed in the methods).

We found that the stepwise decrease of input DNA quantity from optimal 200 ng down to 6.25pg led to a gradual decrease in the SNP call rate (**Figure 2, Table S2**). In particular, with manufacturer-recommended 200 ng, call rates for all 24 individuals were high as expected at 99.9% (SD 0.0%), and decreased to an average of 96.7% (SD 2.5%) with 1 ng, 90.6% (SD 5.0%) with 250 pg, 84.4% (SD 4.5%) with 100 pg, and further down to a minimal call rate of 42.8% (SD 6.3%) from the lowest amount of 6.25 pg. While the number of called SNPs decreased with decreasing input DNA amounts, an increase in the number of false genotypes was seen, as may be expected, while differently so for the different types of genotype errors. Errors were classified as “false heterozygotes” when a homozygous SNP was incorrectly typed as a heterozygote, and “false homozygotes” when a heterozygous SNP was incorrectly typed as a homozygote. The error rate started to increase already with the first DNA dilution step below the manufacturer recommended 200 ng i.e. with 1 ng, albeit with a small average of 1.8% (SD 2.4%) for heterozygote errors, while homozygote errors remained close to zero at 0.01% (SD 0.01%) (**Figure 2, Table S2**). With 250 pg the heterozygote error rate increased to an average of 7.9% (SD 6.8%) and the homozygote error rate to an average of 0.15% (SD 0.27%). Both error rates gradually increased further with further stepwise decreased input DNA amounts, with the heterozygote error rate more so than the homozygote error rate. With 25 pg, where the kinship classification success rate was zero for all degrees of relationship, the heterozygote error rate was on average 43.3% (SD 9.7%), while the homozygote error rate was 8.1% (SD 2.6%). With the lowest input DNA amount of 6.25 pg, the heterozygous and homozygous error rates were highest with on average 72.6% (SD 2.7%) and 21.3% (SD 4.3%), respectively (**Figure 2, Table S2**).

**Figure 2:**
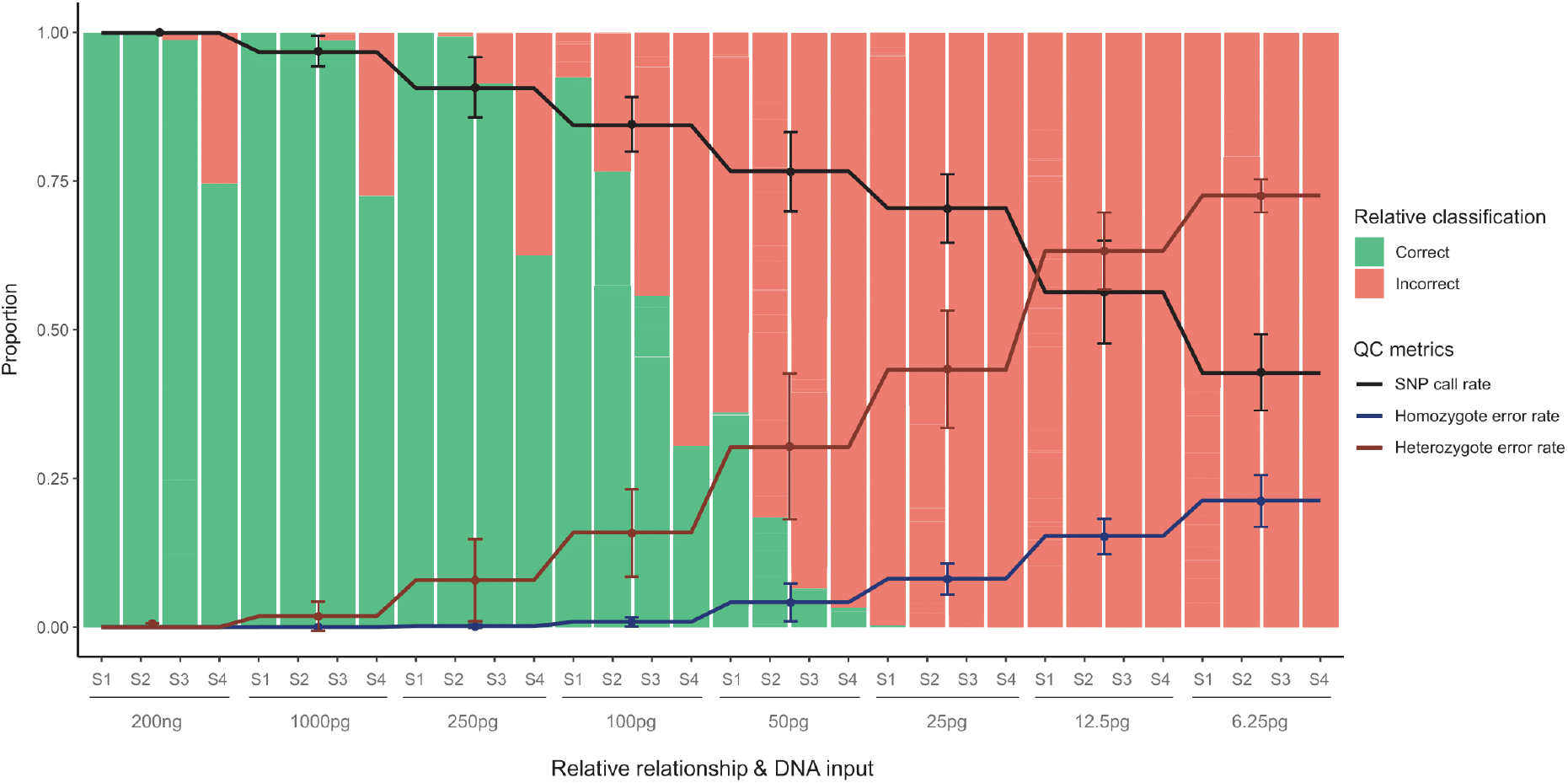
Quality metrics of SNP microarray data from input DNA of decreased quantity. The x-axis represents the 8 stepwise decreased input DNA quantities tested (200 ng – 6.25 pg) and the degrees of kinship classification (S1 - Full siblings, S2 - 1^st^ cousins, S3 - 2^nd^ cousins, S4 - 3^rd^ cousins) across the 24 individuals used. The y-axis runs from 0-100%, depicting call rate, homozygote error rate, heterozygote error rate and classification success. Genotype call rate and error rates are the average of the 24 genotype datasets obtained for each input DNA quantity level. Error bars represent the 75% confidence interval.

### Impact of decreased input DNA quality on SNP microarray-based kinship classification

Further, we investigated the effect of compromised DNA quality on kinship classification success. This included non-degraded DNA samples as recommended by the microarray manufacturer and degraded samples in three DNA fragmentation steps with decreased average fragment size of 1000 bp, 500 bp and 150 bp for 12 individuals, of which 1 ng of DNA was used for SNP microarray analysis (details in the methods section). Data from sample #526 at the fragmentation level of 1000 bp was removed from the final data set for being an outlier (Supplementary Table 9).

We observed that a stepwise decrease of input DNA quality by increasing the severity of DNA fragmentation led to a gradual decrease in kinship classification success (**Figure 3, Table S3**). More specifically, while an average fragment size of 1,000 bp (least severe degradation level tested) had no effect on the classification success for siblings for all 12 individuals (100% success), the first cousin classification success of two individuals was no longer 100% (average success of 99.8%). A more severe effect was seen for the classification of 2^nd^ cousins, where success decreased to an average of 89.7% from the 98.3% obtained with non-fragmented DNA. An average fragment size of 500 bp resulted in a significant drop of kinship classification success for all degrees of relatives tested i.e., to 56.7% for siblings (5 samples (41.7%) still at 100% success), 45.8% for 1^st^ cousins (1 sample still at 100% success, while 4 samples were at 0% success), and 19% for 2^nd^ cousins (5 samples at 0% success rate). As also seen in the DNA quantification experiments for optimal DNA amounts, for 3^rd^ cousins the classification success was already reduced with non-degraded DNA (average 70.8%), and decreased further to 53.7% at 1000 bp fragmentation, and further to 9.5% at 500 bp. Finally, the genotype profiles obtained from the most severely fragmented DNA samples of 150 bp were insufficient to perform any accurate kinship classification (0%) for any of the relatives tested. Similar to the DNA quantification experiments, there were no cases where compromised input DNA quality resulted in an increase in the degree of kinship misclassification (no false positives); in fact, all samples not correctly classified only saw a decrease in the degree of kinship misclassification (**Figure 3, Table S3**).

**Figure 3:**
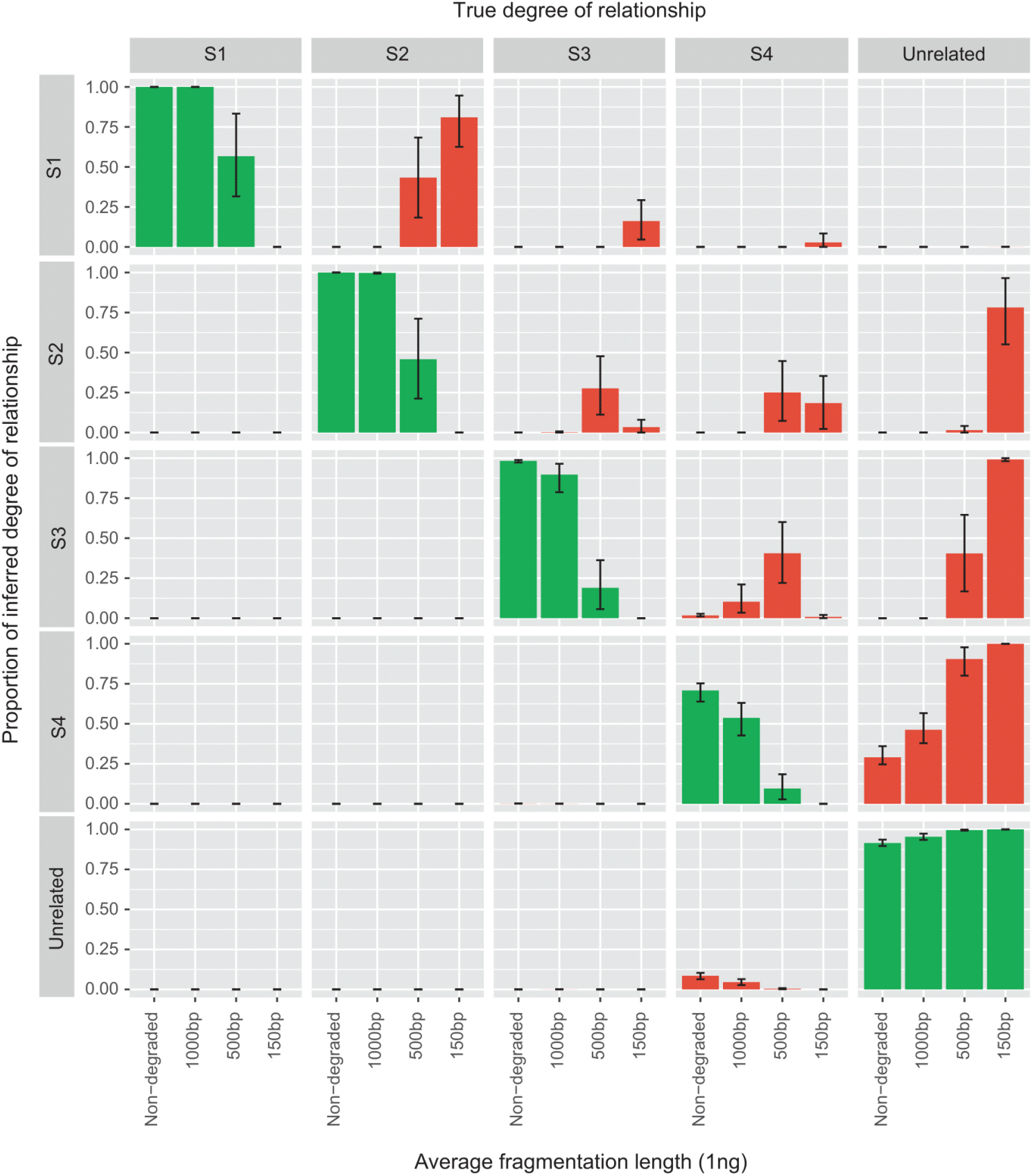
Kinship classification of simulated relatives based on SNP microarray data from input DNA of decreased quality. Each plot in this matrix represents a simulated kinship relation and a specific kinship classification based on a SNP microarray dataset obtained from decreasing input DNA fragment size. Each of the 12 samples used in this experiment had 1000 relatives simulated for each degree of relative, which were S1 (Full siblings), S2 (1^st^ cousins), S3 (2^nd^ cousins), S4 (3^rd^ cousins) or unrelated from the high-quality reference data generated from manufacturer-recommended input-DNA quantity and quality; the kinship classification was also restricted to these possible relation categories. For each of these 12 individuals, four input DNA fragmentation levels i.e., unfragmented, 100 bp, 500 bp, 150 bp were processed on the GSA to obtain a genetic dataset. The estimated genetic relation between such quality-compromised dataset and the relatives simulated from high quality reference data was then classified for each of the 12 individuals. The x-axis of the matrix describes the classified relations, while the y-axis of the matrix describes the original simulated relations, across the 12 individuals used. For the individual plots, the x-axis describes the average fragment size of the input DNA, and the y-axis describes the average kinship classification rate. The green bar plots represent correctly classified relation and the red bar plots represent the incorrect classifications. Error bars display the 75% confidence interval.

### Effect of decreased input DNA quality on SNP microarray genotype accuracy

Aiming to better understand the observed negative impact decreased DNA quality has on kinship classification success, we studied the effect of quality-compromised input DNA on SNP call rate and genotype error rate. Stepwise decrease of DNA quality by increase of DNA fragmentation from non-fragmented down to highly fragmented resulted in a gradual decrease in SNP call cate and an increase in genotyping errors, as may be expected. In particular, an average fragment size of 1,000 bp had an immediate negative effect on the SNP call rate with an average of 84.6% (SD 4.4%) and a negative effect on genotype accuracy with increasing heterozygous error of 16.6% (SD 6.5%) on average and homozygous error of 0.2% (SD 0.2%), compared to the non-degraded DNA samples with averages of 96.5% (SD 1.7%) 1.7% (SD 1.3%) and 0.0% (SD 0.0%), respectively. An average fragment size of 500 bp resulted in a further decrease of the SNP call rate to an average of 75.1% (SD 5.7%), and a further decrease of the genotype accuracy with increasing average heterozygote error of 32.4% (SD 10.0%) and 2.3% (SD 1.5%) homozygote error. Finally, the most severely fragmented input DNA of 150 bp had an average SNP call rate of 62.7% (SD 7.2%), an average heterozygote error of 53.6% (SD 9.7%), and an average homozygote error of 8.1% (SD 2.7%) (**Figure 4, Table S4**).

**Figure 4:**
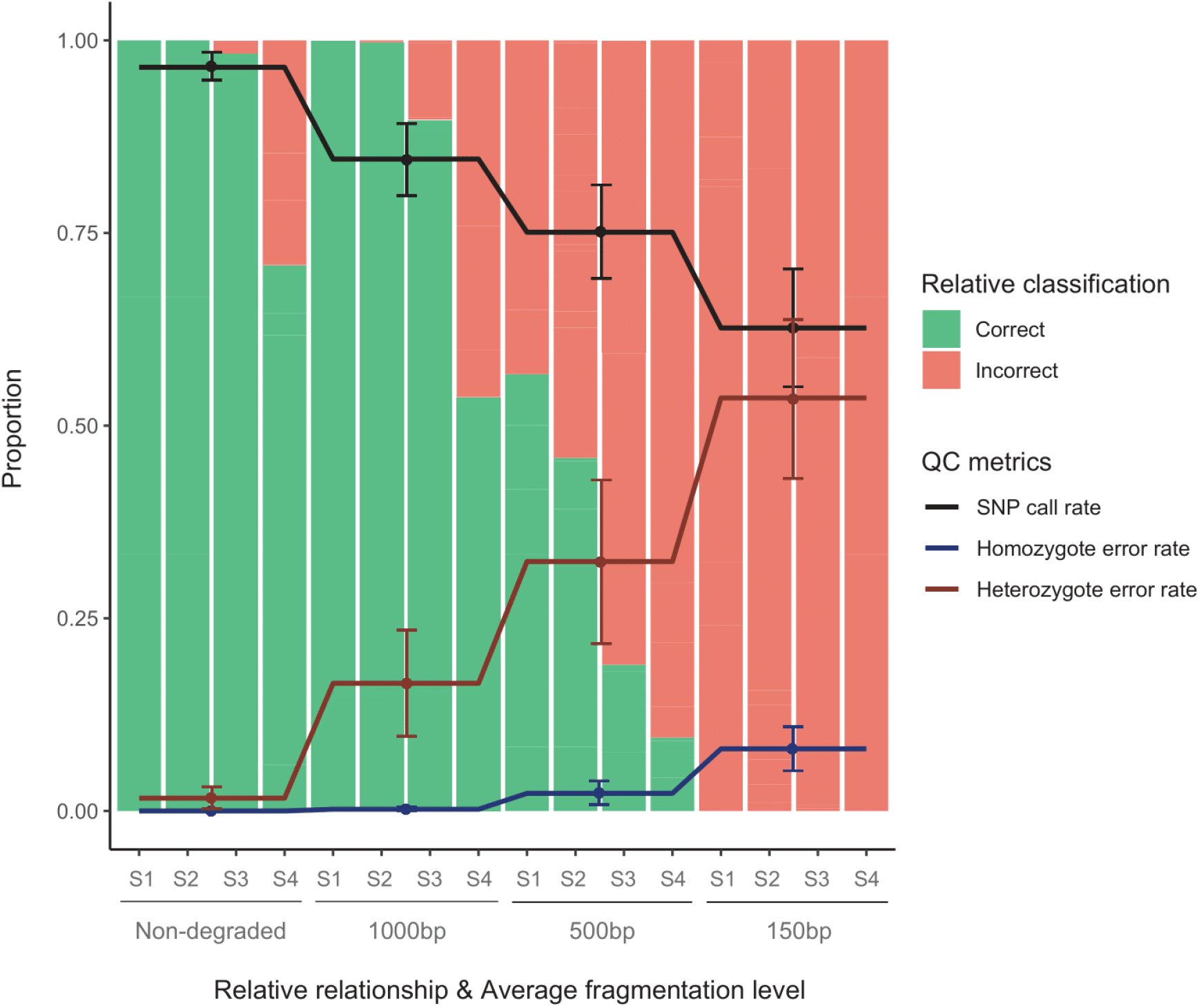
Quality metrics of the GSA genotype data from input DNA of decreased quality. The x-axis represents the 4 stepwise fragmentation levels tested (unfragmented, 1000bp, 500bp and 150bp) and the degrees of kinship classification (S1 - Full siblings, S2 - 1^st^ cousins, S3 - 2^nd^ cousins, S4 - 3^rd^ cousins) across the 12 individuals used. The y-axis runs from 0-100%, depicting call rate, homozygote error rate, heterozygote error rate and classification success. Genotype call rate and error rates are the average of the 12 genotype datasets obtained for each input DNA fragmentation level. Error bars represent the 75% confidence interval.

### Additional factors influencing SNP microarray genotype accuracy and SNP microarray-based kinship classification

To further test SNP microarray performance on the type of DNA samples typically confronted with in forensic casework, and its impact on kinship classification success in the context of IGG, we performed SNP microarray analysis of forensic mock casework DNA samples. Mock casework samples are a typical element of forensic validation studies; they are produced in a way to mimic real crime scene samples. These mock casework samples were produced from blood of individuals for which we had generated high-quality reference data, thus allowing to estimate genotype accuracy and testing the impact on kinship classification. To this end, we ran a total of 55 mock casework DNA samples from i) one individual’s whole blood in making 24 bloodstains of different size (blood volume), prepared on different substrates, and exposed under different environmental conditions such as storage time, temperature, humidity, and UV radiation, the latter to mimic sun exposure, ii) blood DNA samples from three individuals at 200 ng on eight different artificial PCR inhibition levels, and iii) seven non-human DNA samples (for details see material and methods) (**Table S5**).

Overall, we found that varying levels of blood stain storage conditions regarding humidity, substrate type and storage time (up to 25 days) did not seem to have any effect on the SNP microarray genotype accuracy and thus not on the kinship classification success (**Table S5**). In contrast, and as expected, the size of bloodstain (DNA quantity) and DNA damage via direct UV treatment (DNA quality) both appeared to affect genotype accuracy and kinship classification success (**Table S5**). Particularly for the smallest bloodstains produced from 1 µl of blood, the total isolated DNA amounts ranged from 0.167 to 5.163 ng depending on various conditions. The effect of low DNA amount was evident in bloodstain 17 (296 pg), which resulted in a lower accuracy of classification of 3^rd^ cousins (47.1%), in line with what we expected from the DNA quantification experiments (**Figure 1**, **Table S1)**. However, when we combined the effect of low DNA quantity with low DNA quality by damaging small amounts of DNA with direct UV treatment (30 minutes) as in bloodstain 22 (167 pg), we observed a decreased classification accuracy for 2^nd^ cousins of 52.2% compared to 89.7% in the non-treated DNA sample (**Table S5**). Notably, this decrease of kinship classification success was not evident in bloodstain 23 despite the increased time of UV exposure (60 minutes), likely caused by the higher input DNA amount (1.35 ng) in this sample (**Table S5**). Finally, our chosen PCR inhibitor, hematin (27), did not seem to affect SNP microarray genotype accuracy. Independently from the hematin concentration, all samples yielded perfect call rates of over 99% (**Table S6**). Given the very high call rate observed, estimation of kinship classification success was not performed with the SNP array data from the hematin experiment. Moreover, SNP microarray genotyping of all seven animal cell DNA samples, derived from common pets and domesticated animals whose DNA is often recovered from crime scenes, resulted in very low SNP call rates below 70% (**Table S7**), highlighting the human specificity of the GSA. Notably, such low call rates were in our systematic DNA quantity and quality experiments only observed for the ‘worst’ human DNA samples, namely those with lowest input DNA quantity (12.5 and 6.25 pg) (**Figure 2**) and quality (150 bp fragment size) (**Figure 4**).

### Correlation between QC metrics from SNP microarray data and SNP microarray-based kinship classification

In typical practical applications of SNP microarray genotyping, such as for IGG but also for other forensic and non-forensic purposes, only the SNP call rate is available as quality control (QC) measure. Nevertheless, users of SNP microarray technology in practical applications shall be interested in the reliability of the data they obtained from their analyzed DNA samples. As a first step towards providing guidance on this matter of high practical relevance, we tested if our diverse set of experimentally generated SNP microarray data from various input DNA quantities and qualities would allow us to establish a preliminary measure on the reliability of SNP microarray when deviating from the manufacturer recommendations for input DNA quantity and quality. To this end, aiming at maximizing the statistical power, we pooled all data we generated from high and low quality and quantity human DNA samples i.e., from DNA quantity, DNA quality, and mock human bloodstain experiments, resulting in a combined dataset containing call rate, genotype errors rates and classification successes from a total of 264 DNA samples **(Table S5, Table S8, Table S9)**. Using these data, and by employing n-parameter logistic regression to fit a model, we correlated total SNP call rate with i) total genotype error rate, ii) heterozygote error rate and iii) homozygote error rate and obtained very high positive r^2^ estimates of 0.98, 0.99, and 0.95, respectively (**Figure 5a**). Furthermore, we correlated total SNP call rate with kinship classification success rate for all four degrees of relatives (**Figure 5b**), where the fitted model achieved an r^2^ of 0.89 for siblings, 0.87 for 1^st^ cousins, 0.95 for 2^nd^ cousins, and 0.94 for 3^rd^ cousins. We also made a model correlating homozygote error rate with kinship classification success and achieved an r^2^ of 1.00 for siblings, 1.00 for 1^st^ cousins, 1.00 for 2^nd^ cousins, and 0.98 for 3^rd^ cousins (**Figure S2**).

**Figure 5:**
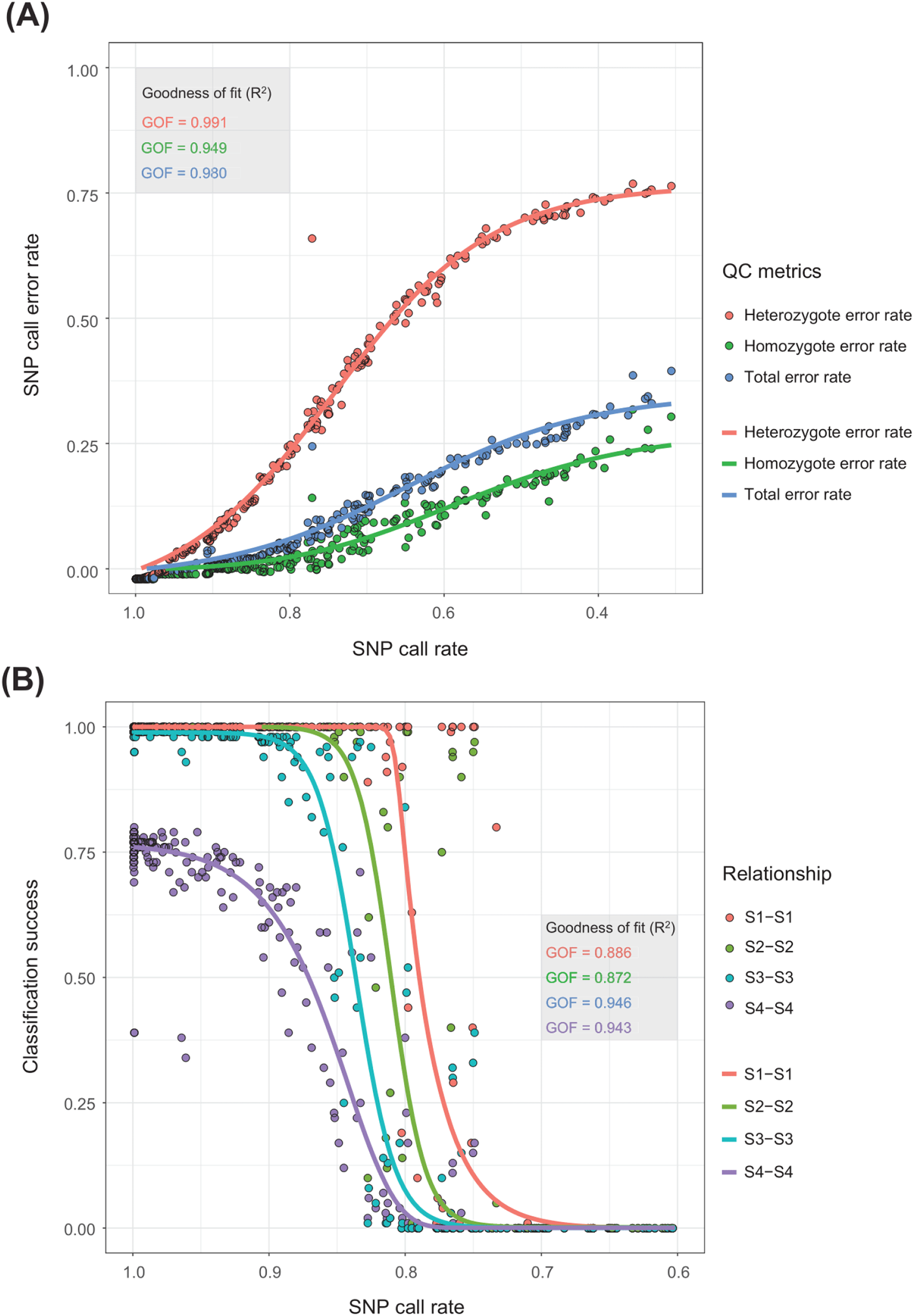
Relations of SNP microarray data quality metrics and kinship classification. To describe the relation between the studied quality control metrics of our experiment, models were fitted with n-parameter logistic regression, using the genotype error, call rate and kinship classification success data from the compromised quantity and quality as well as mock casework SNP microarray analysis in a total of 264 samples (Table S5, Table S8, Table S9). **A)** Relationship of SNP call rate and genotyping error rate. Three types of genotyping errors were considered for this model: homozygote error, heterozygote error, and the total genotyping error by combining both. The homozygote error model had a standard error (SE) of 0.016 with a goodness-of-fit (GoF / r^2^) of 0.948, the heterozygote error model had a SE of 0.0247 with a GoF of 0.991, and the total error model a SE of 0.014 with a GoF of 0.980. **B)** Relationship of SNP call rate and kinship classification success. One model was fitted for each degree of relation: S1-S1 (full sibling), S2-S2 (1^st^ cousin), S3-S3 (2^nd^ cousin), and S4-S4 (3^rd^ cousin). The full sibling classification success prediction model had a SE of 0.163 with a GoF of 0.886. The 1^st^ cousin success prediction model had a SE of 0.174 with a GoF of 0.872. The 2^nd^ cousin success prediction model had a SE of 0.110 with a GoF of 0.946. The 3^rd^ cousin success prediction model had a SE of 0.084 with a GoF of 0.943.

As may be expected, we saw that all three genotyping error rates (total, heterozygote and homozygote) increased as the SNP call rate decreased, with the heterozygote error being impacted the most (**Figure 5a**). While the heterozygote error rate increased immediately with decreased SNP call rate, the homozygote error rate remained below 0.05% as long as 93% of the array SNPs were called (**Figure 5a**). More importantly, we found that within certain boundaries it was possible to predict kinship classification success solely based on SNP call rate (**Figure 5b**). Whereas for the most distant relationships, i.e., 3^rd^ cousins, kinship classification rate gradually decreased with call rate, the success of classifying siblings, as well as 1^st^ and 2^nd^ cousins, remained at 100% until the call rate reached certain thresholds. These thresholds were different for the different degrees of relationship and were lower the more closely related the family members were. With SNP call rates below these thresholds, kinship classification success decreased rapidly (**Figure 5b).** More specifically, we observed a remaining high kinship classification success until, and drastic decrease below, 90% SNP call rate for 2^nd^ cousins, ∼86% for 1^st^ cousins, and ∼82% for siblings. Universally and independently of family relationship, we found that DNA samples with SNP call rates below 75% appeared unsuitable for correctly inferring kinship relations, as below this threshold we observed 0% classification success for all four degrees of relationship.

### SNP microarray analysis of naturally degraded DNA from skeletal remains

Finally, to assess SNP microarray performance on naturally degraded DNA, thereby combining compromised DNA quantity with compromised DNA quality through a natural body decomposition process, we analyzed DNA samples obtained from skeletal remains buried underground or found on forest surface that had experienced various natural conditions for various times of postmortem intervals. A total of 18 bone or teeth derived DNA samples were GSA-genotyped (**Table S10**). These DNA samples were additionally genotyped with a commercial forensic STR kit commonly used for forensic identification. The latter allows a comparison between the performance of both DNA technologies. Due to the nature of this sampling set-up, SNP microarray reference data obtained from manufacturer-recommended input DNA quantity and quality were not available for the individuals of which bone DNA samples were analyzed, thus not allowing genotype accuracy estimations and kinship classification success quantifications. However, by applying our established relation between SNP call rate and kinship classification success (see above, **Figure 5b**), we used the SNP call rate obtained from these naturally degraded DNA samples as proxy for SNP microarray data quality, and from the obtained results, provided expectations on kinship classification success in case such samples would be used for IGG purposes.

Two out of the 18 bone DNA samples (bones 17 & 18) with the highest recovered DNA amounts of 27.09 and 78.5 ng fully consumed for GSA analysis, produced microarray-based SNP genotype profiles with a call rate above 95% (**Table S10**). According to our fitted models, these data could be used to correctly classify relatives up to 3^rd^ cousins (**Figure 5b**). These two bone DNA samples also yielded complete 24-loci forensic DNA profiles. Two other bone DNA samples (bones 14 & 16) with total input DNA amounts of 2.16 ng and 3.14 ng yielded call rates of 85.3% and 88.7%, respectively (**Table S10**), which according to our fitted models could still be used to classify relatives up to 3^rd^ cousins (**Figure 5b**), albeit with reduced success rate. Sample 14 yielded a complete 24-locus forensic DNA profile, while sample 16 missed two non-autosomal loci. Additionally, 12 of the bone DNA samples with total input DNA amounts at the picogram level resulted in call rates below 70% (ranging from 32.6% to 69.7%), which according to our models is unsuited to obtain any accurate kinship classification (**Figure 5a**). Of these 12 bone DNA samples, 7 gave complete 24-loci forensic DNA profiles, while 5 had partial profiles between 14 and 22 of the 24 markers. The remaining two bone DNA samples (bones 1 and 2) with total input DNA amounts of 36 pg and 56 pg failed the GSA analysis completely as they produced no signals. In the forensic DNA profiling, these two DNA samples had produced results for 16 and 22 of the 24 loci, respectively.

## Discussion

Our work represents the first study that systematically explores the impact SNP microarray analysis of quantity and quality compromised DNA has for kinship classification success, which is relevant for IGG. Earlier studies on SNP microarray genotyping of forensic DNA samples did not report genotype reliability and whether the genetic data would be applicable for kinship classification (18) or investigated the use of SNP microarrays for low quantities of DNA in less samples and not analyzing DNA below 1 ng (23, 25). Moreover, as far as we are aware, no previous study empirically investigated the impact of quantity and/or quality compromised DNA on SNP microarray-based kinship classification success, which, however, is highly relevant for applying SNP microarrays for IGG purposes in forensic practice.

The model commonly used in genetic genealogy for classifying relatives from genomic data implements versions of the so-called segment approach (13, 28–30), which we also applied here in a modified way. The novelty in our approach lies in the combination of conditional simulation with the final step in which the original (perfect), gold standard reference genotype, is replaced with an imperfect sample (e.g., DNA of low quantity and/or quality). Using this novel approach on the SNP microarray data acquired from optimal manufacturer-recommended input DNA of high quantity and quality, 25% of the 3^rd^ cousins were not correctly classified, while for 2^nd^ cousins this only was 1%, and for 1^st^ cousins and for siblings we had 100% classification success (**Figure 1**). This finding is in line with expectations, as the more distant the family relationship is, the less likely one will be able to detect shared DNA because recombination can make the shared segment shorter or can lead to not inheriting the shared DNA at all (31).

In our quantity-compromised DNA experiments, an amount of 250 pg of high molecular weight DNA yielded similarly high success rates of near 100% for classifying siblings and 1^st^ cousins, and 1 ng for 2^nd^ cousin, as did the manufacturer-recommended 200 ng. Although these reduced input DNA quantities resulted in decreased SNP call rates and increased rates with which homozygous loci are erroneously called as heterozygotes, we did not notice any measurable impact on kinship classification success. This picture changed when the input DNA quantity was decreased below these critical amounts, which resulted in gradually decreased kinship classification success down to 25 pg DNA and lower, where kinship classification success was zero for all four degrees of relatives tested. While an increase in the homozygote error rate was observed, the kinship classification success rates dropped.

On the other hand, in our quality-compromised DNA microarray experiments, when 1 ng of DNA was gradually fragmented and used for SNP microarray analysis, the first fragmentation level of 1000 bp yielded similarly high success rates of 100% for classifying siblings and 1^st^ cousins as the non-fragmented input DNA did, while decreases in SNP call rates and increases in genotype error rate were seen compared to non-fragmented input DNA. For 2^nd^ cousins and 1000 bp fragmentation, a reduced classification success was found, which further decreased for all relatives with further increased fragmentation levels. At the highest fragmentation level of 150 bp, the kinship classification success was zero for all degrees of relatives. These findings together imply the main deciding factor if kinship classification based on SNP microarray data from compromised DNA is successful or not is the amount of proper quality DNA in a sample.

Another novelty of our work is that we empirically determined how GSA microarray genotype data is affected when compromised DNA is analyzed. We found that false heterozygotes were much more prevalent than false homozygotes when degrading the DNA quality and quantity. Previous work has highlighted how detrimental genotyping errors are for relative classification (32) (33). We analyzed both genotyping error types separately as these were different in prevalence and how they affected relative classification rate. Our data strongly suggests that the false homozygote genotypes are the driving cause for relative misclassification. This is expected, as the segment approach is highly sensitive to homozygote genotype errors. Only a false homozygote can cause IBS0 (where neither of the alleles are similar between the individuals) and pre-maturely end a shared segment. This is corroborated by our finding that homozygote genotyping error rate and kinship classification success rates are strongly correlated (**r^2^** of 0.999 for siblings, **r^2^** of 1.00 for 1^st^ cousins, **r^2^** of 0.999 for 2^nd^ cousins and a **r^2^** of 0.976 for 3^rd^ cousins) (**Figure S2**). The false homozygote genotype calls will terminate segments in the kinship classification model rather than prolonging them; thus, resulting in a decreased degree of inferred relatedness instead of an increased one. This could explain why decreasing DNA quantity or quality did not inflate the false positive kinship classification rate i.e., the degree of relatedness was always underestimated in our experiments.

Other models to infer family relationships from genomic data may have a different sensitivity to genotyping errors, which may be explored in future studies. In particular, the likelihood ratio-based model, adopted in forensic STR profiling, could be an alternative (32, 34, 35). With this model, fewer markers are needed in the calculations compared to the segment model (32). Nevertheless, a virtue of the segment approach used here is that it is unaffected by the most frequent error of Illumina GSA SNP genotyping microarrays as we showed here i.e., heterozygote genotyping errors. Another advantage of the segment approach is that it allows kinship classification to be estimated in the absence of specifically stated pairs of hypotheses of relatedness, which instead is a key requirement for applying the likelihood ratio approach. We note that several implementations of the segment approach mitigate issues with genotyping errors, for instance through allowing a number of opposite homozygote genotypes in a shared segment (29, 36).

As a side aspect, we studied the isolated effect of decreasing SNP numbers, as proxy for SNP call rate, on the kinship classification success under optimal input DNA conditions. For this, the used the high-quality reference data, leaving aside the effects of decreased input DNA quantity and quality. This exercise illustrates that for closer relationships (e.g., siblings) SNP genotyping call rates as low as 5.6% can still allow accurately classification through detection of shared segments, while for more distant relationships (e.g. 3^rd^ cousins) it needs SNP genotyping call rates above 50% to be sufficient (**Figure S3**). These minimal SNP numbers rather serve as theoretical expectations, whereas practically relevant thresholds of SNP call rates to achieve trustworthy kinship classifications are seen in Figure 5. These findings confirm our observations from quantity and quality compromised DNA samples that SNP genotyping call rate alone is not driving kinship classification success (**Figures 2, 4 & 5**). This further corroborates the finding that SNP genotyping errors are the main driving force of kinship misclassification, even so, a decrease in SNP genotyping call rate can be accounted for in the segment approach by tuning the number of SNPs required to define IBD segments. However, we demonstrated that decreasing this threshold can have a detrimental impact on kinship classification success (**Figure S4)**. Decreasing the number of SNPs per segment will lead to false inclusion of random segments as IBD, thus, resulting in an estimated higher degree of family relationship compared to the true degree.

Remarkably, we had fairly high kinship classification success with low input DNA amounts of 100 pg. We attribute this to the whole genome amplification (WGA) step that is part of the standard SNP microarray protocol. WGA increases the amount of DNA several thousand fold (37, 38), potentially compensating in part for a low input DNA amount. WGA is typically not utilized in forensic DNA analysis, as it can cause STR profiling to result in extraneous bands (39) and unequal amplification between different targeted loci (40). While these concerns are relevant for forensic STR profiling, the detrimental effect of WGA on SNP microarray genotypes is reported to be considerably lower (38, 41). WGA-based SNP microarray genotype data were found to be highly concordant with the SNP microarray data from the respective gDNA (42). An additional benefit of WGA is that it is unaffected by the presence of the PCR-inhibitor hematin (**Table S6**). However, the assumed WGA-based compensation effect of low input DNA quantity has limits, as our study clearly demonstrated.

By decreasing the DNA quantity and quality below the recommendations of the microarray manufacturer, we also observed a notable rise of genotyping errors, where the increase of false heterozygous genotypes was larger than that of false homozygote genotypes (**Figures 2 & 4**). The intrinsic mechanisms of the Illumina GSA suggest an explanation why heterozygote genotype calls are more prone to be false than homozygote genotype calls. We provide a brief description of Infinium technique behind Illumina GSA arrays in **Figure S5**. We hypothesize that the majority of false genotype calls observed with decreased input DNA quantity and quality was caused by decreasing probe intensity due to less DNA being bound to hybridization probes, and a subsequent effect on the signal processing and genotype interpretation software **(Figures S6 & S6)**. When probe intensity decreases, the chance increases that the SNP call shifts to a different genotyping cluster. The heterozygote cluster is found closest to the lowest intensity values, which may cause homozygotes to be called as heterozygotes, explaining the higher heterozygote error rate compared to homozygote error rate, which we observed at increasingly compromised input DNA for both input DNA quantity and quality.

Judging the suitability of a DNA sample for SNP microarray analysis is generally difficult since a number of factors such as quantity, fragmentation, and DNA damage could impact genotype accuracy. In typical practical applications of SNP microarrays to compromised DNA, such as IGG, reference data from manufacturer-recommended input DNA quantity and quality are unavailable so that genotyping errors cannot be determined. Therefore, we used quality metrics that are available for every SNP microarray dataset, such as SNP call rate, to address if a SNP microarray dataset is suitable for kinship classification, or not. Our results show that SNP genotyping call rate is highly correlated with genotyping error rate and with kinship classification success rate (**Figure 5a&b)**. Therefore, we suggest that the SNP call rate, being the most common SNP microarray genotyping quality metric available, can be used as a primary indicator to estimate the reliability of Illumina GSA genotype datasets, and thus its suitability for kinship classification such as in IGG. In our data analysis, we observed a steady increase in false SNP genotypes in SNP microarray datasets with SNP call rates below 99% (**Figure 5b**). Different SNP genotyping call rate thresholds are applied by different users, with 95% being one standard call rate cut-off (43, 44). We found that samples with SNP call rates of 90% or above yielded reliable kinship classification; while below 90% we started to observe a decrease in kinship classification success, first for 2^nd^ cousins, then on lower call rate levels also for 1^st^ cousins and for siblings (**Figure 5b)**. Third cousin classification started at a 75% success rate even at high quality and quantity input DNA and declined gradually with decreasing call rates. Based on our data, for SNP genotyping call rates below 75% it is unlikely that kinship classification will be successful for any degree of family relationship. Consequently, there is a grey zone between 75% and 90% call rate where the kinship classification success is still high for close relationships (e.g., siblings), but quickly drops for distant relationships (e.g., 2^nd^ cousins). Notably, these thresholds coincide with the increase in homozygote error rate, starting at roughly 90% call rate as well (**Figure 5b**).

Since experimental DNA degradation is artificial and does not resemble natural degradation processes, we additionally analyzed bone and tooth derived DNA samples from skeletal remains. These DNA samples naturally degraded under varying conditions and combine decreased DNA quantity and decreased quality, which is typical in missing person cases (45). However, only a minority of the skeleton-derived naturally degraded DNA samples yielded high-enough call rates sufficient for concluding high genotype accuracy (two samples had call rate > 93%, two others were in the grey zone >85% call rate). These four bone DNA samples also had comparatively high amounts of quantified DNA, highlighting the importance of input DNA amount for GSA analysis. The qPCR-measured input DNA amounts of these four best performing bone DNA samples were all above 2.1 ng, while we obtained similar call rates of ∼85% using 100pg of non-fragmented DNA. As our DNA quality experiments have shown that severely degraded DNA cannot be genotyped accurately with the GSA, the lower SNP microarray genotyping reliability can be attributed to the poor quality of nuclear DNA in the majority of the bone or teeth DNA samples tested (45). Notably, the SNP microarray results of these bone and teeth derived DNA samples were not in good agreement with results from forensic STR profiling. Although we found that 3 of the 4 samples with high SNP call rates had complete forensic STR profiles, this was also seen for 7 of the samples with low SNP call rates. A difference in success rates of forensic STR profiling and SNP microarrays is not unexpected given the substantial differences in the underlying genotyping technologies and in the number of analyzed DNA markers with each technology.

While the majority of naturally degraded DNA samples from skeletal remains did not result in reliable SNP genotype profiles, our mock casework samples that simulated crime scene samples generally did result in high quality genotyping profiles as well as high kinship classification success (**Table S5**). This is most likely due to the fact that blood stains generally are a good source for DNA (46) and that the artificial DNA degradation applied to the mock casework samples was less severe in impact and time than the natural DNA degradation in the skeletal remains. These preliminary data provide optimistic expectations that SNP microarray genotyping might find application in the forensic cases, including IGG, as long as DNA yield is high enough e.g., with input DNA amounts above 250-500 pg, and DNA degradation is rather low, such as obtained from crime scene stains that experienced rather mild environmental conditions similar to what we simulated in our mock casework samples. For DNA samples of lower quantities and/or more severe degradation, as seen in the skeletal remain samples and the severely fragmented DNA samples analyzed here, SNP microarray analysis is rather not suitable. Besides DNA quantification to select samples for SNP microarray analysis, the achieved SNP call rate and the preliminary models we introduced here can be used as guidance on the suitability of the generated data such as for kinship classification in IGG applications.

While our study is the first to empirically investigated the impact of quantity and/or quality compromised DNA on SNP microarray-based kinship classification success in the context of IGG, others previously reported on the use of SNP microarrays on forensic DNA samples, or other samples with quantity limitations. The company Parabon Nanolabs (USA) reported results from about 250 forensic case samples genotyped on the CytoSNP-850k array (18), a predecessor of the Illumina GSA array (47). Roughly half of the DNA samples analyzed with SNP microarrays were reported with SNP call rates above 95%, while also about half had more than 10 ng of input DNA used (18). Based on our analyses, those cases are expected to find kinship matches in IGG database investigations using the segment approach as long as the respective relative is included in the database used. Wendt et al. (23) recently performed DNA titration experiments from 200 ng to 1 ng on 3 DNA samples obtained from postmortem blood samples with Illumina array (Omni2.5Exome-8 v1.3) genotyping as part of their DNA input pretesting for a genome-wide association study (GWAS). They found that the SNP microarray genotypes from 200 ng down to 25 ng did not differ significantly and thus used 25 ng as input DNA threshold for their GWAS. Similar to our findings, which were based on larger sample size and larger titration span going down to 6.25 pg, the authors showed higher rates of allelic drop-in (false heterozygotes) than allelic drop-out (false homozygotes) with decreasing DNA quantity. Another recently published study explored how long-time stored serum samples, with an average input DNA amount of 5.8 ng, performed on two other types of SNP microarrays, the Affymetrix Axiom Array (Thermofisher) and the Illumina HumanCoreExome array. In 80% of these DNA samples, SNP call rates above 94% were reported. In comparison, 92% of our 1 ng high-quality DNA samples had SNP call rates above 94%. The level of degradation was not quantified in that previous study, which could account for the lower number of reported samples with high call rate. However, the authors had made alterations to the standard SNP microarray genotyping protocol to account for the expected low DNA quality, given the use of long-time stored serum. For Axiom Array analysis the number of WGA cycles was doubled, and for the Illumina Human Core Exome array analysis DNA restoration kits were used (25). These are options that might improve SNP call rates, which we did not explore here and may deserve systematic investigation in the future.

In conclusion, our study provides the first empirical evidence how SNP microarray analysis of quantity- and quality-compromised DNA impacts on kinship classification success, which is highly relevant in the context of IGG. The GSA used here as an example of a widely used SNP microarray was able to obtain high-density SNP profiles that were accurate enough to achieve high kinship classification success at 800 times lower input DNA amounts for siblings and 1^st^ cousins, and 200 times lower for 2^nd^ cousins, compared to the amount recommended by the microarray manufacturer. Decreasing DNA quantity or quality reduced the number of called SNPs and introduced heterozygote and, to a lesser extent, homozygote genotyping errors. These SNP microarray genotyping errors, believed to be caused by low probe intensity, are the primary cause of incorrect kinship classification as result of using compromised DNA for SNP microarray analysis. When applied for IGG purposes, the consequence may potentially be to miss a match to a relative included in the genetic genealogy database, while the genotyping errors are unlikely to result in a false positive kinship match as our study implies. Overall, our results are relevant for SNP microarray applications of compromised DNA for IGG purposes, aiming at finding unknown perpetrators of crime via their relatives stored in genomic databases accessible by law enforcement agencies. They are also relevant for other forensic SNP microarray applications such as appearance prediction and biogeographic ancestry inference of unknown perpetrators or missing persons from compromised DNA, aiming at helping to identify unknown perpetrators and missing persons via focused investigative intelligence.

## Materials and Methods

### Samples

Two types of human biological samples were used in this study: i) high-quality EDTA blood samples from 24 completely anonymous blood donors of European descent that A.U.G. received in 2003 from the Sanquin blood bank in Rotterdam, Netherlands, for the purpose of genetic research and genetic method evaluation, and ii) DNA samples from 18 skeletons collected by a forensic pathologist after medico-legal examination of human remains found on a forest surface or exhumed for genetic identification, at the Department of Forensic Medicine in Krakow, Poland, including teeth, skull bones, humeri, femoral, clavicle and metacarpal bone. Sample details and description of DNA isolation and quality control can be found in **Supplementary Methods**.

### DNA titration

The manufacturer-recommended amount of input DNA for the Illumina GSA microarray, used here as a prominent SNP microarray example, is 200 ng (48). This is beyond what is typically available in forensic crime scene samples where the DNA yield is highly various and depends on the size of the crime scene stain and other parameters. In this series of experiments, we therefore tested for the effect of the input DNA amount below the optimal 200 ng on the SNP microarray performance in terms of genotyping suboptimal amounts of DNA i.e. 1000 pg, 250 pg, 100 pg, 50 pg, 25 pg, 12.5 pg and 6.25 pg, produced via DNA titration and confirmed via a qPCR assay (Qiagen Investigator Quantiplex Kit). The minimal amount of 6.25 pg roughly equals the genomic DNA content of one human diploid cell. Given that for each of the 24 individuals tested we analyzed 8 different input DNA amounts, this DNA quantity experiment included a total of 192 DNA samples that were processed for SNP microarray genotyping.

### DNA fragmentation

The manufacturer’s recommendation on input DNA quality for the Illumina GSA microarray is to use high molecular weight DNA. DNA samples obtained from crime scene stains are usually of low molecular weight due to DNA degradation caused by various factors that impact on biological stains at the scene of crime, such as temperature and humidity, or long time periods since sample deposition at the scene of crime prior to collection. In this series of experiments, we therefore tested for the effect of input DNA with decreased quality. We opted for DNA fragmentation as one component of DNA degradation and used the adaptive focused acoustics technology (Covaris), which has the advantage of fragmenting DNA to a preset average fragment length. For 12 samples, randomly sampled from the initial 24, we degraded 1 ng of DNA to average fragment sizes of 1,000 bp, 500 bp, and 150 bp as analyzed on the Labchip GX (Perkin Elmer). Given that for each of the 12 individuals tested we analyzed 4 different input DNA amounts, this DNA quality experiment included a total of 48 DNA samples that were processed for SNP microarray genotyping.

### Mock casework

To test for the performance of the Illumina GSA microarray on forensic-type samples, we generated mock casework samples that mimic crime scene stains. In total, 24 bloodstains from whole blood of one of the 24 blood donors were exposed to different conditions that are often encountered in forensic samples. The factors we considered included stain size (the volume of blood used), storage time, storage temperature, substrate (the surface blood was deposited), relative humidity and DNA damage (via UV radiation), as detailed in **Table S5**. The total amount of DNA extracted from each bloodstain was used as input in the Illumina GSA array, ranging from 0.167 to 127.8 ng of DNA. Additionally, we tested the effects of PCR inhibitor hematin and non-human DNA on GSA genotyping, detailed in the supplementary methods.

### Microarray genotyping

The study used the Infinium Global Screening Array (GSA) for all SNP microarray genotyping (48). The choice was made partly based on its popularity in the Direct-to-Consumer Genetics industry, but also due to its rich up-to-date SNP content. Array genotyping was performed according to standard Illumina protocols (37). We decided to refrain from additional sample or data treatment, e.g. the use of restoration kits on the DNA samples or application of Hardy-Weinberg equilibrium on the SNP data. Microarray scan data was converted to genotypes using Genome Studio 2.0, the standard software for processing Illumina SNP microarrays. As this project consisted of samples of sub-optimal input DNA quantity and quality, standard microarray QC protocols, such as filtering on SNP or sample call rates, were not executed. We extracted a subset of markers based on the overlap with the 1000 Genomes phase III reference panel (49), resulting in a total of 519,300 SNP markers. Genetic positions in centiMorgan (cM) were extracted from Rutger’s map or alternatively interpolated for markers absent from the repository (50). For each gold standard sample, the genotypes were phased to generate haplotypes employing the Eagle V2.4 algorithm (51) with the European (CEU) individuals from the 1000 Genomes Project as reference data (49).

### Genotype quality assessment

The high-density SNP profiles obtained from GSA analysis of each tested condition was rated by two main measures of quality: SNP call rate and genotype discordance to the gold standard, which was obtained from manufacturer-recommended input DNA quantity and quality for each of the 24 individuals. SNP call rate is the fraction of the total probes with genotype calls and it is the most common quality measure used in SNP microarray analysis. The number of false genotypes in each dataset was determined by direct comparison of the genetic profile from each tested condition to their corresponding gold standard reference dataset obtained from optimal DNA conditions. Heterozygous error rate is defined as the percentage of false heterozygotes among all heterozygotes, and homozygous error rate is defined as the percentage of homozygotes that are either heterozygotes or opposite homozygote in the gold standard reference data.

### Conditional simulations of relatives

For each of the 24 individuals, we generated relatives through a conditional simulation approach using the gold standard reference dataset obtained from manufacturer-recommended input DNA quantity and quality. **Figure S1** provides an overview of our algorithm to simulate family members based on our gold standard samples. In brief, the approach uses phased genotype data from high quality samples and proceeds by conditionally generating relatives for each sample. We restricted the algorithm to siblings, 1^st^ cousins, 2^nd^ cousins and 3^rd^ cousins (In the figures abbreviated as S1, S2, S3 and S4), spanning a range of the most relevant degrees of relationships for IGG purposes. For each of the 24 gold standard samples, we repeated this relative simulation process 1000 times for each type of relation. In detail, the simulation process starts at the first SNP on a chromosome and initiates by drawing alleles from the gold standard sample with probabilities equal to the identical by descent (IBD) probabilities for each degree of relationship (14, 52). Moving along the chromosome, the process either continues with the same IBD state as for the previous marker or change IBD state with probabilities equal to the rate of recombination between the markers. Ultimately, the process generates a complete high-density SNP dataset for the relative with a mosaic of shared DNA with the gold standard sample. In contrast to approaches that employ population allele frequencies and gene dropping to generate pedigree data, our approach i) takes into account the known phased genotype of the original sample and ii) uses phased haplotypes from the population to draw non-IBD alleles; thus, mitigating potential biases caused by linkage disequilibrium. The novelty in our approach is the combination of conditional simulation with the final step in which the original, gold standard, genotype is replaced with an imperfect sample.

### Inferring degree of relatedness

Genealogy assessment, referred here as kinship classification, was performed by inferring the degree of relatedness of the imperfect sample to the simulated relatives (see Figure S1). The term *imperfect* is used as an umbrella for inhibited, degraded or quantity-reduced DNA samples. We used a version of what we call the segment approach to infer degree of relatedness between individuals. The algorithm is described in for example (13, 29, 30) and detailed for forensic purposes in Kling et al. (14). The version implemented by direct-to-consumer company Ancestry.com is also outlined in a white paper (28). This is a method that identifies haplotypes between individuals that are identical by descent (IBD) without requiring information on allele frequencies in the population. Briefly, the most naïve version of the approach measures stretches of genotypes where at least one nucleotide is identical at each base position between the pair of individuals (IBS1 or IBS2) to ultimately infer IBD segments. The stretches are only terminated if opposite homozygous genotypes are detected (IBS0). Furthermore, in order to accurately define a segment as IBD the algorithm needs thresholds, firstly the length as measured in cM and secondly the total number of overlapping SNPs in each segment. Both thresholds have the purpose of excluding non-IBD segments. The length restriction was tuned to include short enough segments for distant relatives and still large enough not to include unrelated individuals as relatives. We used 5 cM as the length detection threshold, corroborated by previous studies (53) as well as our own studies (**Figure S7**). We further explored the impact of the second threshold i.e., the number of overlapping SNPs required in each segment. As the imperfect samples may contain several locus dropouts, a default requirement of say 500 SNPs can result in missed segments. Therefore, we explored 100-700 SNPs as thresholds, detailed in **Figure S6**. Finally, in order to explore the isolated effect of decreasing number of markers, we thinned the data, resulting in reduced marker subsets ranging from the original 519,300 down to 10,379 markers (**Figure S3**). We proceeded to infer relatedness with each reduced set of markers using the methods previously described.

## Author Contributions

J.H.V., D.K., A.V., A.G.U., M.K. designed study with contributions from T.J.P.;

J.H.V., D.K., A.V., P.A., V.K., M.M.P.J.V. generated data;

J.H.V., D.K., A.V. analyzed data;

A.V. prepared main display items with contributions from J.H.V. and D.K.;

A.G.U., D.P.-R., M.K. contributed resources;

J.H.V., D.K., A.V., M.K. wrote the manuscript;

P.A., V.K., M.M.P.J.V., D.P.-R., T.J.P., A.G.U. edited versions of the manuscript;

All authors approved the final manuscript.

## Acknowledgments

The work of J.H.V., P.A., M.V., A.G.U., A.V., V.K., M.K. was supported by Erasmus MC, University Medical Center Rotterdam. AV was additionally supported with an EUR Fellowship by Erasmus University Rotterdam. We thank Benjamin Planterose Jiménez (Department of Genetic Identification, Erasmus MC University Medical Center Rotterdam) for providing technical assistance and suggestions in implementing R, and Wojciech Branicki (Malopolska Centre of Biotechnology, Jagiellonian University) for his assistance with making bone DNA samples available to this study.

## Supporting information

### Supplementary Methods

#### Supplementary Figures

Figure S1. Description of our novel approach to conditionally generate relatives

Figure S2. Homozygote error rate *versus* family classification success

Figure S3. Isolated effect of call rate on family classification success.

Figure S4. Effect of segment SNP number and total call rates on family classification success.

Figure S5. Simplified technical background of Illumina GSA.

Figure S6. Possible explanation fot the high number of false heterozygotes in compromised DNA genotyping.

Figure S7. Effect of segment length on family classification success.

#### Supplementary Tables

Table S1. Proportion of inferred relationship for all degrees of relationship of the titration samples.

Table S2. SNP microarray QC metrics of the titration samples

Table S3. Proportion of inferred relationship for all degrees of relationship of the fragmented samples

Table S4. SNP microarray QC metrics of the fragmented samples

Table S5. Overview of the mock casework samples including QC metrics (call and genotype discordance rates) and family classification success

Table S6. Overview of artificially inhibited samples and their call rates

Table S7. Overview of animal DNA samples and their call rates

Table S8: Individual sample data for the quantity experiment

Table S9: Individual sample data of the quality (DNA fragmentation) experiment

Table S10. Information on the samples obtained from skeletal remains included in this study

