## Supplementary Figures for "Impact of SNP microarray analysis of compromised DNA on kinship classification success in the context of investigative genetic genealogy"

**This PDF file includes:**

Figures S1 to S7

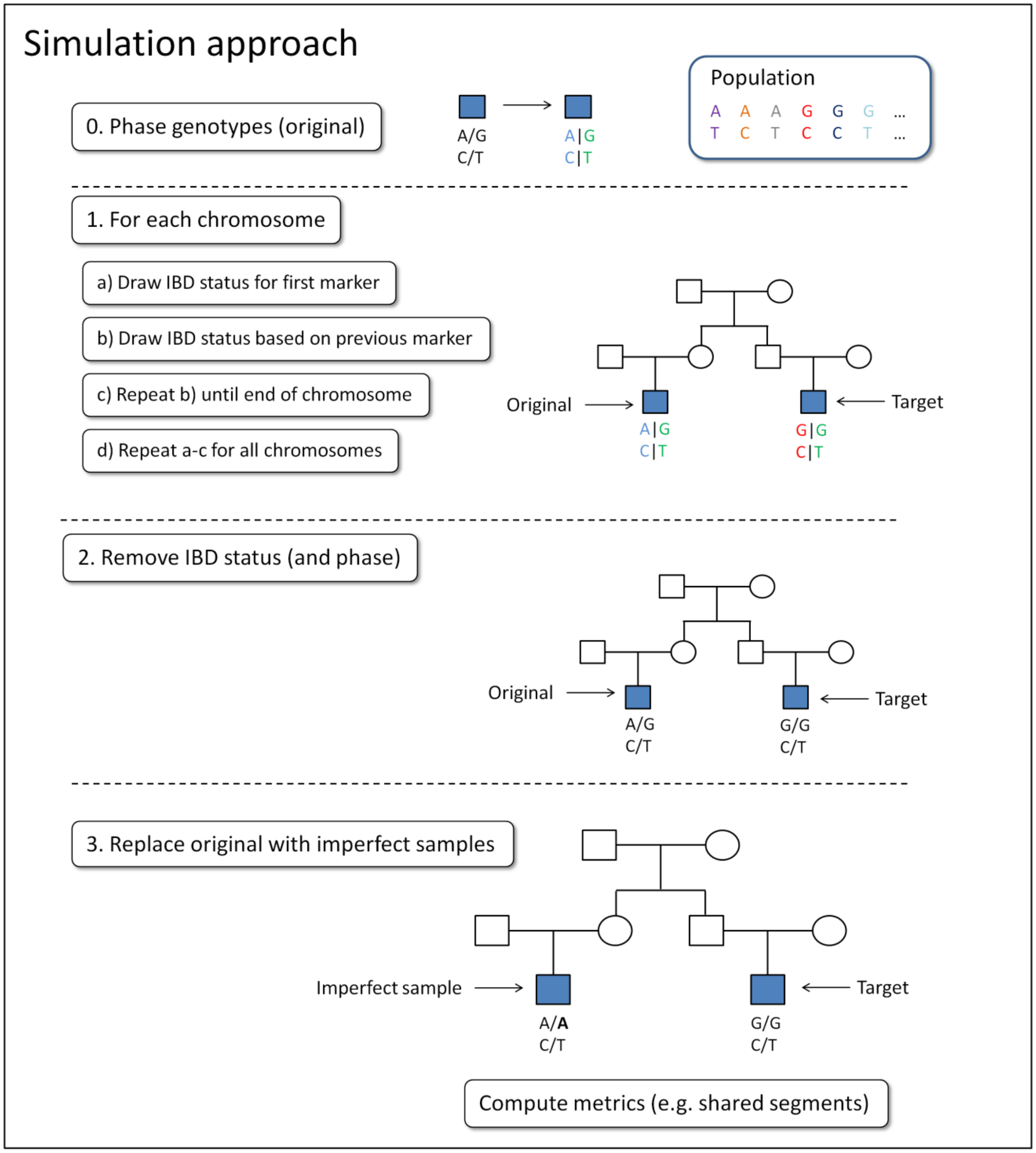

Simulation approach
0. Phase genotypes (original)
A/G
C/T
A|G
C|T
Population
A A A G G G ...
T C T C C T ...
1. For each chromosome
a) Draw IBD status for first marker
b) Draw IBD status based on previous marker
c) Repeat b) until end of chromosome
d) Repeat a-c for all chromosomes
Original
Target
A|G
C|T
G|G
C|T
2. Remove IBD status (and phase)
Original
Target
A/G
C/T
G/G
C/T
3. Replace original with imperfect samples
Imperfect sample
Target
A/A
C/T
G/G
C/T
Compute metrics (e.g. shared segments)

**Fig. S1. Description of our novel approach to conditionally generate relatives (illustrated for two SNPs).** The algorithm starts with a genotype with unknown phase, denoted 'Original'. The genotype is first phased as described in the main text. Our simulation starts using the phased genotypes of for the original sample (also referred to as *Golden standard*) For each chromosome the algorithm randomly draws the genotype of a relative, denoted 'Target' (illustrated as a first cousin in the figure). The identical by descent (IBD) probabilities for first cousins compose Pr(IBD=0)=0.75, Pr(IBD=1)=0.25 and Pr(IBD=2)=0. The novelty is illustrated in the final step of our algorithm whereby the original (Golden standard) sample is replaced with an imperfect sample, e.g. a mock or low quality DNA sample. Ultimately, we compute relationship metrics using the target (relative) and the imperfect sample in addition to performing computations using the original sample (for comparative reasons).

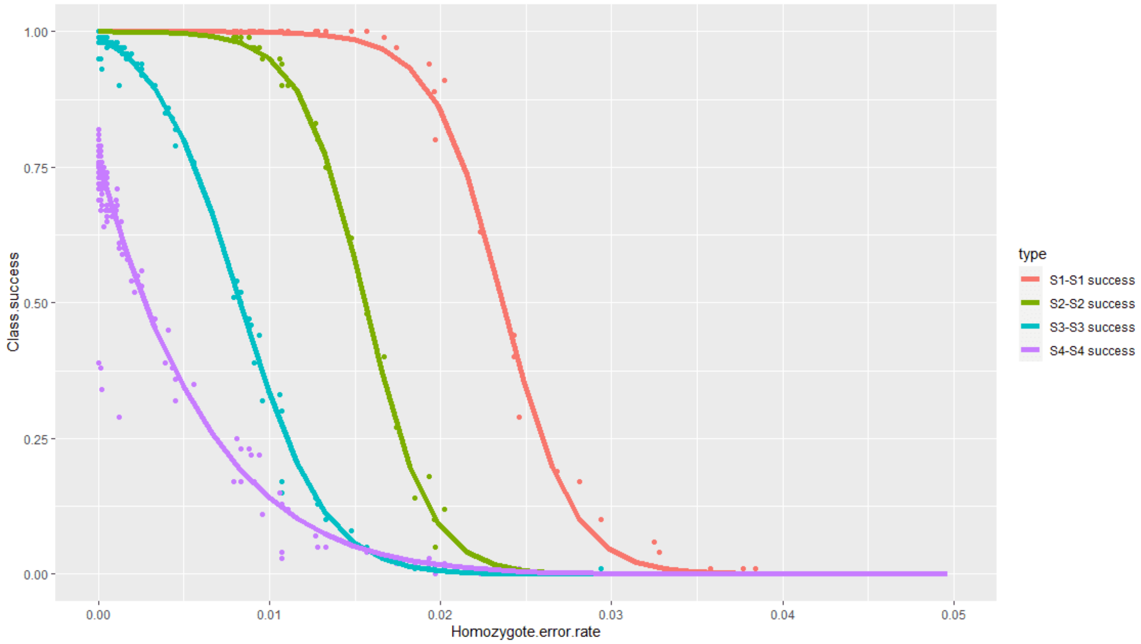


**Fig. S2**. **Homozygote error rate *versus* family classification success**. Models were fitted with N-parameter logistic regression for each degree of relationship. The S1-S1 success prediction model has a standard error (SE) of 0.011 with a $r^2$ of 0.999.The S2-S2 success prediction model has a SE of 0.008 with a $r^2$ of 1.00. The S3-S3 success prediction model has a SE of 0.114 with a $r^2$ of 0.999. The S4-S4 success prediction model has a SE of 0.055 with a $r^2$ of 0.976. S1: Full siblings, S2: First cousins, S3: Second cousins and S4: Third cousins.

**Text for figures S3 and S4:**

**Impact of SNP number on microarray-based kinship classification for optimal input DNA**

To expand on the isolated effect of call rate, we investigated the impact the total number of SNPs has on kinship classification success under optimal input DNA conditions (i.e. manufacturer-recommended 200 ng high-quality DNA). To this end, we iteratively thinned a total number of 519,300 SNPs extracted from the high-quality reference data set from the 24 individuals (cross-referenced with data from the 1000 Genomes Project, see methods for details), and estimated kinship classification rates with decreasing numbers of SNPs, thereby mimicking the effect of decreasing SNP call rates (see methods). We found that with 161,000 SNPs, corresponding to a call rate of 31%, and less, the classification rates of 3rd cousins rapidly started to drop, while successful classification of 2nd cousins started to drop with 105,400 SNPs (20.3% call rate), and that of 1st cousins with 79,000 SNPs (15.2% call rate). Classification of siblings was reliable down to only 31,700 SNPs, corresponding to a call rate of only 6.1% (Figure S3). This shows that distant relationships are generally more sensitive to reduced number of SNPs than closer relationships, as would be expected. However, since these results were obtained from optimal input DNA amounts by using our high-quality reference data, and the thinning process omitted full loci rather than individual alleles with a locus and did not consider any other types of genotyping error, these numbers may not fully mimic the loss of allelic data that results when low quality and/or quantity DNA is tested. In our empirical data from quantity and quality compromised input DNA, only one sample had a call rate below 31%. At the lowest input DNA quantity level of 6.25pg, the SNP call rate was on average 42.8%. At the lowest input DNA quality level of 150 bp fragmentation, the average SNP call rate was 62.7%, with the lowest individual SNP call rate being 46.4% (Figure 2, Figure 4, Table S8, Table S9).

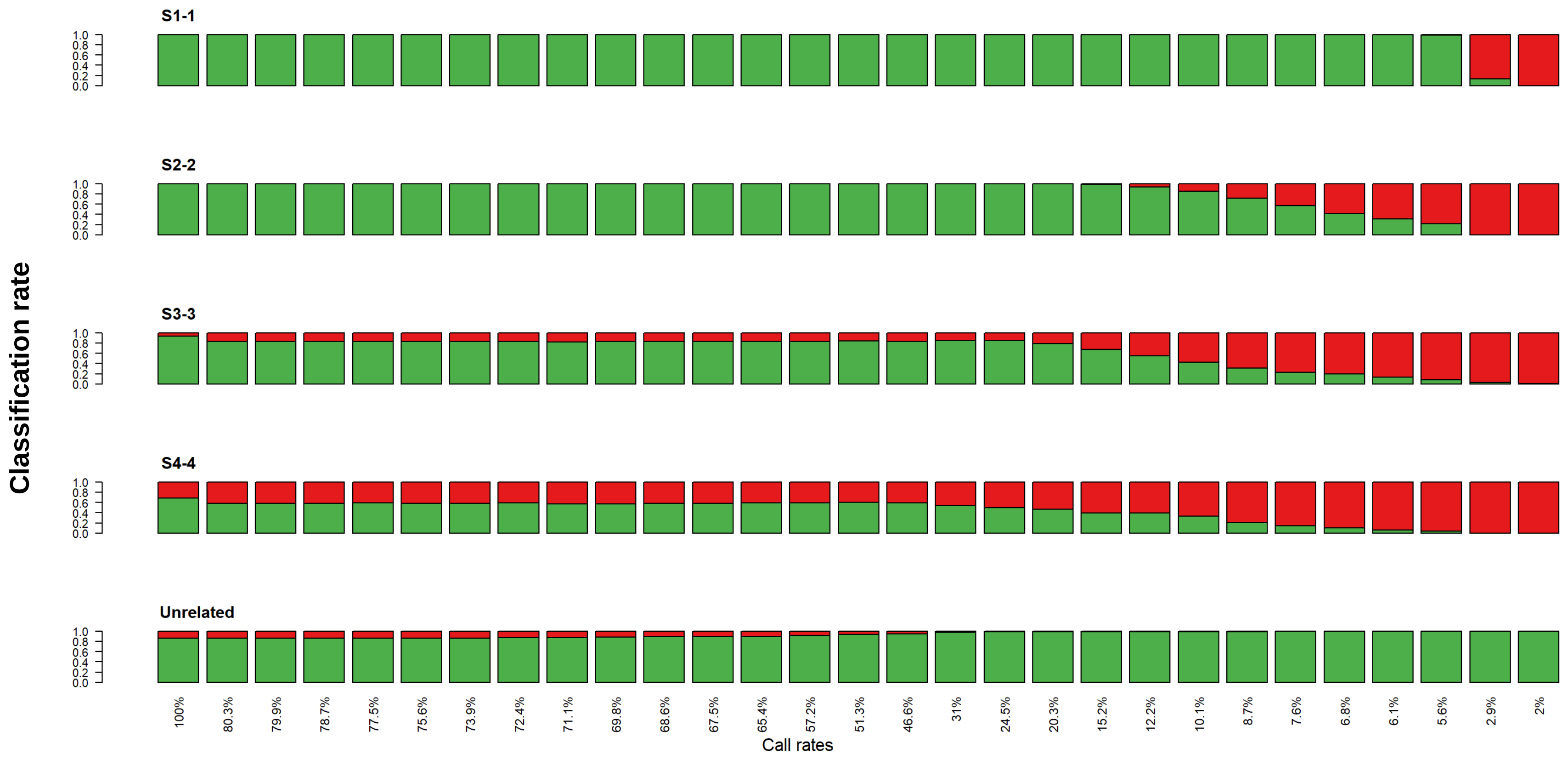


**Fig. S3. Isolated effect of call rate on family classification success.** Family classification success for 1000 simulated pairs of relatives for each degree of relationship based on the complete set of 519,300 SNP markers but on a decreasing call rate. Each bar plot indicates the rate of true (green) and false (red) classifications, without reference to the type of misclassification. The x-axis represents the call rate, ranging from 100% (519,300) to 2% (10,379). The graph illustrates, as expected, that closer relationships require less markers (lower call rates) to provide correct classifications. For instance, a pair of full siblings requires as little as 5-6% (26,000) of the markers to give an almost 100% correct classification rate. While, on the other hand, more distant relationships where the individual identical-by-descent segments are smaller, more and denser genetic marker data is needed. For instance, a pair of third cousins needs at least 50% (259,000) of the markers to provide acceptable classification rates. However, and importantly, call rates above 50%, if the marker data is evenly spread, is generally sufficient for the relationships and methods we use. S1-1: Full siblings, S2-2: First cousins, S3-3: Second cousins and S4-4: Third cousins.

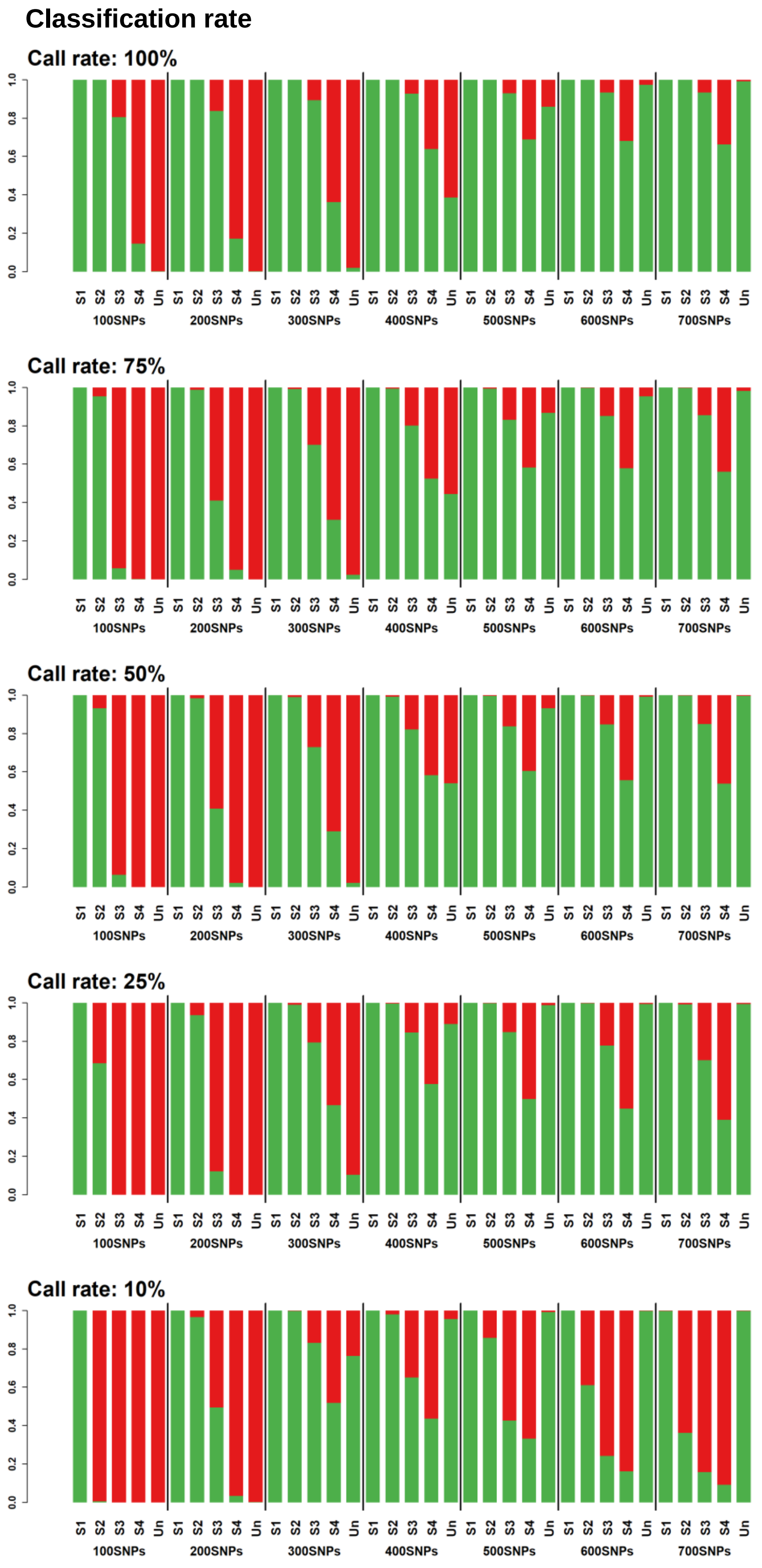
Classification rate
Call rate: 100%
Call rate: 75%
Call rate: 50%
Call rate: 25%
Call rate: 10%
0.0
0.2
0.4
0.6
0.8
1.0
S1 S2 S3 S4 Un
100SNPs
200SNPs
300SNPs
400SNPs
500SNPs
600SNPs
700SNPs

**Fig. S4. Effect of segment SNP number and total call rates on family classification success.** Family classifications for 1000 simulated pairs of relatives for each degree of relationship based on decreasing number of SNPs (call rates), from 100% down to 10%. Each individual plot shows classification success per degree of kinship (S1-1:Full siblings, S2-2:First cousins, S3-3:Second cousins and S4-4:Third cousins) on a decreasing number of SNPs used in a segment. We use 5 cM (see Fig S4) as a cut-off to define an identical-by-descent (IBD) segment length between pairs of simulated relatives and explore the required number of SNPs in each such segment. Each bar plot indicates the rate of true (green) and false (red) classifications, without deference to the type of misclassification. The x-axis represents the SNP threshold used in the segment model, ranging from 100 to 700 SNPs. First of all, based on the regular 100% call rate plot, we selected 500 SNPs as a good trade-off between including and excluding true relatives and unrelated individuals.

The interpretation is two-dimensional; focusing on a single row of the plot (i.e. a call rate), a low value on the SNP threshold generally tend to falsely call IBD segment ultimately resulting in an overestimation of the degree of relatedness, for instance “turning” first cousins into full siblings. Isolating a column instead (i.e. a particular SNP threshold), there is a less obvious interpretation. A higher call rate means the genetic comparison is less susceptible to false matches. In summary, for low call rates (e.g. 10%) lowering the SNP threshold to 400 (instead of 500 used in our study) could be a counter-measure to improve classification success.

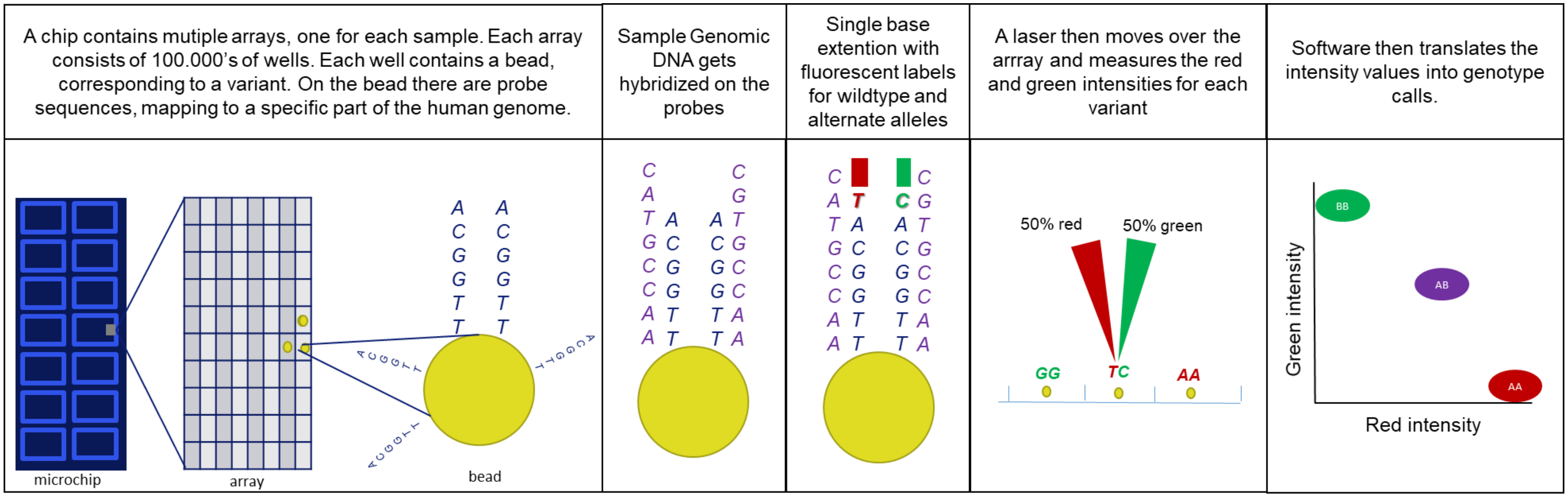


**Fig. S5. Simplified technical background of Illumina GSA. This is a schematic overview of Illumina GSA mechanics of genotype determination. A quick summary of the workings of Illumina GSA array: Each array contains 100.000's of wells containing beads. Each bead has probe sequences on its surface which map to a specific part of the human genome. There are often multiple beads per variant to increase the success chance of correct genotyping. When sample DNA is put on the array, the strands of sample DNA are hybridized onto the matching probes on the array. Then, single base extension at the end of each probe is performed, with two options per bead, often wild type and a variant. One possible allele is attached to the red fluorescent dye and the alternate allele is linked to the green fluorescent dye. When hybridization is complete, lasers will move across the array to measure the intensity of the two colors for each well. This intensity information is then used to determine the genotype in the genotyping software. There the red and green intensities get plotted to determine the genotype. Hence, heterozygous variants have both red and green intensity, while homozygous SNPs deliver intensity from only a single color.**

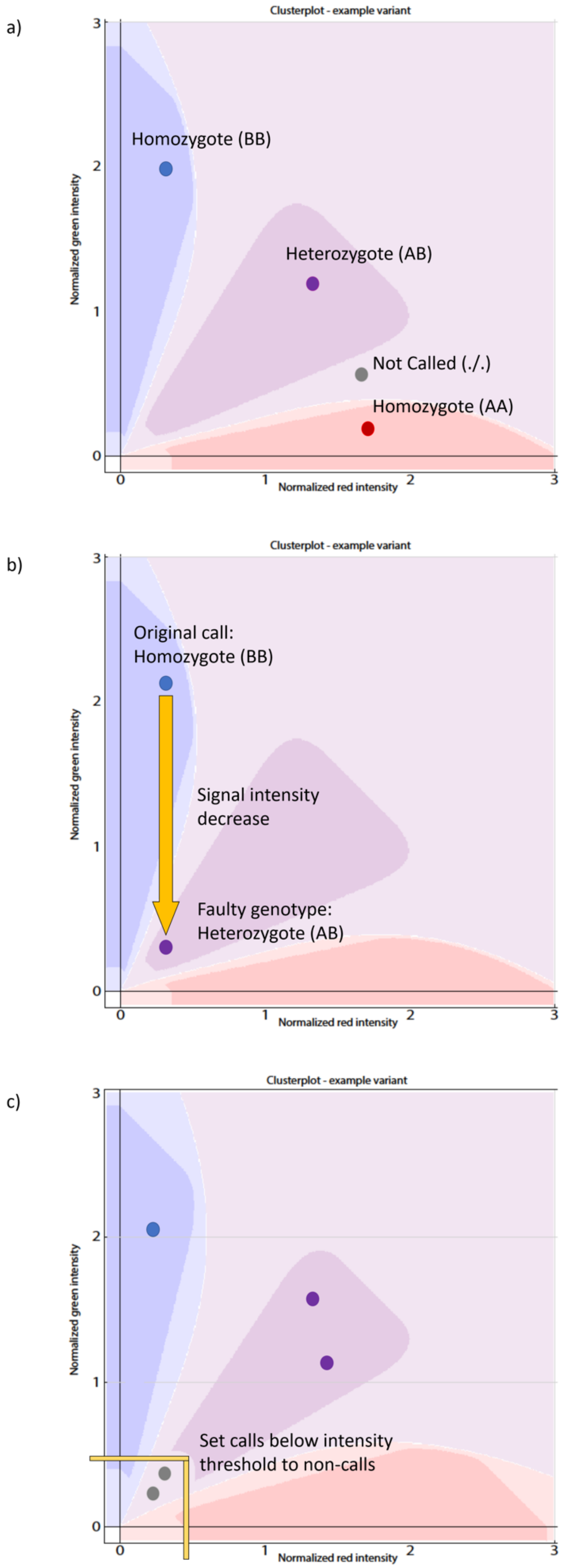
a)
Clusterplot - example variant
Homozygote (BB)
Heterozygote (AB)
Not Called (./.)
Homozygote (AA)
Normalized green intensity
Normalized red intensity
b)
Clusterplot - example variant
Original call:
Homozygote (BB)
Signal intensity
decrease
Faulty genotype:
Heterozygote (AB)
Normalized green intensity
Normalized red intensity
c)
Clusterplot - example variant
Set calls below intensity
threshold to non-calls
Normalized green intensity
Normalized red intensity

**Fig. S6. Possible explanation fot the high number of false heterozygotes in compromised DNA genotyping. This is a followup on the schematic overview of Illumina™ GSA mechanics of genotype determination in figure S4. This is a possible explanation on how wrong genotype calls can be obtained in low-quality samples and why these are mainly heterozygotes.** a) Representation of a SNP probe genotype cluster plot of the Illumina™ GSA. The green and red fluorescent signal of each probe are plotted for each SNP-probe. The intensities of the red and green signal determine in which cluster the signal is found, which determines the genotype call. The boundaries of dye intensities for the genotype calls are determined on beforehand using cluster files. When a call falls outside of the clusters, it is computed as a non-call with no genotype information. b) If the intensity of the signal decreases, a SNP call can occur in the wrong genotype cluster, causing a faulty genotype to be displayed. c) To counter this, an option is to set an intensity threshold for SNP calls. Calls below the set threshold were should be set to non-calls.

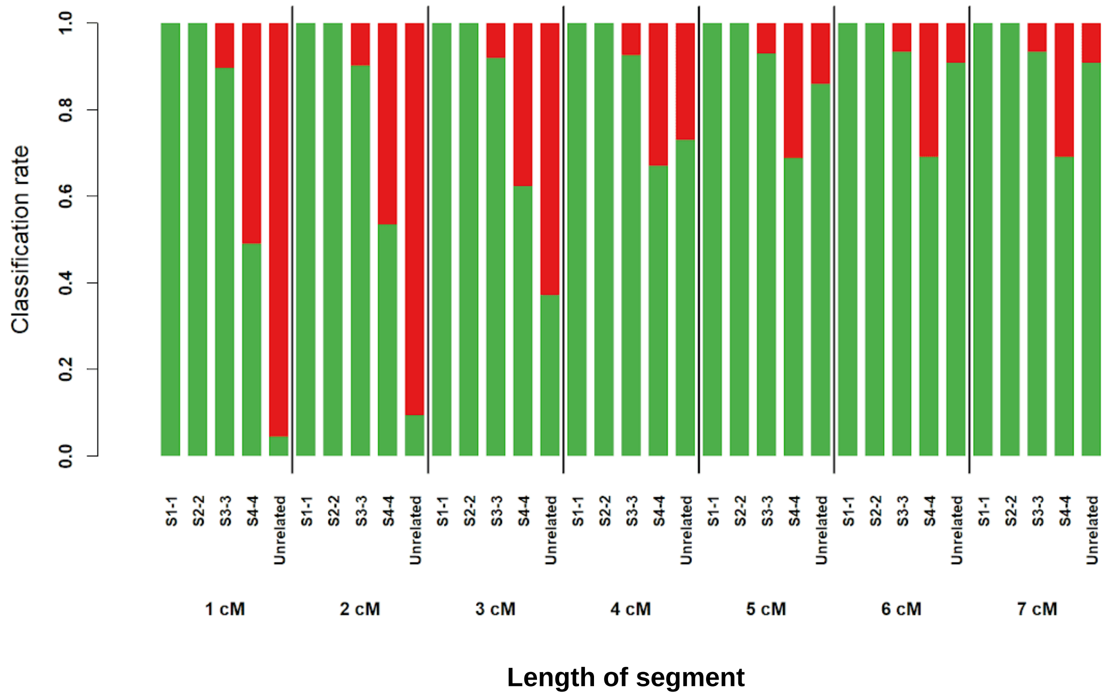


**Fig. S7. Effect of segment length on family classification success.** Family classifications for 1000 simulated pairs of relatives for each degree of relationship based on the complete set of 519,300 SNP markers but on a decreasing length of individual segments that are defined as identical by decent (IBD) in our algorithm. Each bar plot indicates the rate of true (green) and false (red) classifications, without deference to the type of misclassification. The x-axis represents the length threshold (in cM) used in the segment model ranging from 1 to 7 cM. Based on these results, we selected 5 cM as a inclusion threshold, providing a good trade-off to include true relatives and unrelated individuals while simultaneously excluding false relatives. cM: centiMorgan, S1-1: Full siblings, S2-2: First cousins, S3-3: Second cousins and S4-4: Third cousins.
