## Supplementary Methods for "Impact of SNP microarray analysis of compromised DNA on kinship classification success in the context of investigative genetic genealogy"

**This PDF file includes:**

Supplementary methods

### Supplementary Methods

***Blood samples.*** EDTA blood samples were provided by the Sanquin blood bank in Rotterdam in 2003 for the purpose of scientific research and method testing. Genomic DNA was extracted with standard salting out procedures from 5 µl of whole blood. The quality of each extracted DNA sample was assessed by measuring the OD260/280 and OD260/230 ratios with the Nanodrop spectrophotometer (ThermoFisher Scientific, United States). DNA integrity was assessed by running a 0.7% agarose gel. DNA concentrations were initially measured with Nanodrop and the picogreen fluorescence assay (Invitrogen, United States). For more accurate measurements, DNA concentrations were also measured in triplicate with the Investigator Quantiplex kit (Qiagen, Germany) on the Quantstudio 7 Flex (Applied Biosystems, United States). Only DNA samples with an optimal quality (OD260/280: 1.8-1.9, OD260/230: >1.9 and DNA integrity: >10kbp) were selected for the mock case experiments.

***Skeletal remain samples.*** A set of 18 samples, including teeth, skull bones, humeri, femoral, clavicle and metacarpal bone, were collected by a forensic pathologist after medico-legal examination of human remains found on a forest surface or exhumed for genetic identification, at the Department of Forensic Medicine in Krakow, Poland. For remains found on a forest surface post-mortem interval (PMI) was not available, but analyzing effect of environmental conditions and skeletonization of corpse, definitely exceeded one year. The post-mortem interval of exhumed cadavers ranged from 6,5 years to 7,5 years, except one, which was 67 years. The man was arrested and shot in 1951, buried in a nameless grave and exhumed in 2018. The storage time prior to medico-legal examination and bone extraction ranged from 5 to 27 months at -18°C. Genomic DNA isolation was performed up to one week after sample collection. Firstly, bones were treated with 15% bleach, then repeatedly shaken with 70% ethanol and distilled water ($dH_2O$), and finally, UV irradiated for 5 minutes on each side in a UVP CL-1000 Ultraviolet Crosslinker. After decontamination, samples were individually crushed and decalcified using EDTA buffer (0.5M, pH 8.0). DNA was isolated using the Sherlock AX Kit (A&A Biotechnology, Poland) and quantified using the Investigator Quantiplex Kit (Qiagen) on a 7500 Real–time PCR System (Applied Biosystems) according to the manufacturer's instructions. Total DNA amounts from the different samples ranged from 36pg to 78,5ng (**Table S8**). These amounts were fully consumed for GSA genotyping. All DNA samples were genotyped using the GlobalFiler PCR amplification kit (ThermoFisher Scientific) that targets 24 loci, including 21 autosomal short tandem repeats (STRs), 1 Y-STR, and two loci, the X- and Y-copy, of amelogenin. Therefore, females are expected to have a 22-loci complete profile, whereas males a 24-loci profile.

***Mock casework samples***. To test the forensic applicability of the Illumina™ GSA-MD V2 microarrays, we simulated dried mock casework bloodstains under a wide range of forensically relevant conditions, often encountered in forensic casework that could potentially affect downstream analysis. More specifically, a total of 23 bloodstains were prepared using whole blood of a single individual from the EDTA blood samples. The factors that we varied include the volume (µl) of bloodstains, substrate (surface or material that the bloodstain was deposited on), storage temperature, storage time and relative humidity during storage, as well as UV radiation strengths and incubation time. **Table S5** depicts the conditions under which all bloodstains were prepared table sand exposed. Genomic DNA was isolated using the body fluid stains protocol of the QIAamp® DNA Investigator kit (Qiagen) according to manufacturer's instructions. DNA was quantified using the Quantifiler™ Human DNA Quantification kit (Qiagen) on a CFX96 Touch Real-time PCR Detection System (Bio-Rad, United States). Total DNA amounts from these mock bloodstains ranged from 170pg to 222ng . When possible the optimal DNA amount for the Illumina™ GSA was used (200ng). In all other cases the maximum DNA amount (**Table S5**) was used as input to the array.

***Hematin inhibition.*** PCR inhibitors, such as hematin, are commonly found in forensically relevant DNA samples including blood and can negative affect forensic DNA genotyping (Sidstedt et al, 2019, the impact of common PCR inhibitors on forensic MPS analysis). To test whether hematin acts as an inhibitor during the whole-genome amplification included in the Illumina™ GSA analysis, we prepared a total of 24 samples using high-quality DNA (200ng) from three individuals (samples 102,114 and 87, from the original 24 samples) on eight different hematin concentrations (1600uM, 1200uM, 800uM, 400uM, 200uM, 100uM, 50uM, 25uM in 10µl sample volume).

***Animal DNA.*** To test the specificity of the Illumina™ GSA we analyzed DNA from common pets and other animals including mouse, rat, cow, cat, dog, pig and horse. Animal cells were obtained from the Coriell Institute for Medical Research. Genomic DNA was isolated with the DNAdvance kit (Beckman Coulter, United States) and quantified with the picogreen fluorescence assay (Invitrogen). Each animal DNA sample was analyzed on the Illumina™ GSA v2 according to the standard protocol.

***Calibration of the segment approach***. To investigate our approach used to classify relatives, some additional analyses were conducted. Our study used what we refer to as the segment model, whereby the length of shared DNA segments is accumulated across two genetic profiles and subsequently used to determine the degree of relationship. In detail, the model uses half-identical stretches of DNA, such that only one allele needs to be shared between the two profiles. In other words, only opposite homozygote genotypes can terminate a shared segment (given that bi-allelic SNP markers are used). The model has two important input parameters, the length required for a segment to be defined as identical by descent IBD (measured in cM) and the number of SNPs in each called segment. Since appropriately setting these thresholds is crucial for identifying true IBD segments, we conducted simulations to find the optimal settings, in addition to using results from previous studies. More specifically, we varied the number of SNPs in each segment from 100 to 700 (**Figure S4**) and the length of the segment from 1 to 7 cM (**Figure S5**). Additionally, we studied how decreasing the call rate from 100% to 2%, without introducing errors, affects family classifications. In detail, we started with the original set of 519,300 SNPs and for each thinning we varied the inter-marker distance (measured in cM) to decide whether to include a marker in each particular set. The approach creates a genetic map with markers evenly located across the genome (spaced with the given distance threshold). We further performed another set of simulations where we generated 1000 relatives with total call rates 100%, 75%, 50%, 25% and 10% (including errors). We subsequently used the segment model where we varied the threshold on the number of SNPs in an IBD segment from 100 to 700 in order to investigate potential improvements in classification rates
